# rtcisE2F drives liver TIC self-renewal and metastasis via m^6^A-modulated mRNA stability of *E2F6* and *E2F3*

**DOI:** 10.1101/2021.01.24.428027

**Authors:** Zhenzhen Chen, Benyu Liu, Lan Huang, Xiang Zhong, Zhongyi Yan, Pingping Zhu

**Author notes:** These authors contribute equally to this work (Zhenzhen Chen, Benyu Liu, Lan Huang). Corresponding authors: Zhenzhen Chen and Pingping Zhu.

## Abstract

**Background:** Liver tumor initiating cells (TICs) harbor self-renewal and differentiation capacities, and well contribute to liver tumorigenesis, metastasis and heterogeneity. However, the molecular mechanisms of liver TIC self-renewal are unclear. N^6^-methyladenosine is the most abundant modification of RNA molecules, and is involved in RNA stability and translation, but the molecular mechanisms of m^6^A regulation remain largely unknown.

**Methods:** circRNA expression was detected by *in situ* hybridization, fluorescence *in situ* hybridization, quantitative real-time PCR and Northern blot. Target gene expression was examined by microarray analyses, quantitative real-time PCR and Western blot. CRISPR, CRISPR interference (CRISPRi) and short-hairpin RNA (shRNA) were used for circRNA/target gene knockout and knockdown. Liver TICs were enriched through sphere formation and FACS using CD133 as a marker, and liver TIC activity was assessed by tumor propagation, sphere formation, tumor-initiating, and transwell assays. Quantitative real-time PCR and Northern blot were used to determine mRNA stability. RNA–protein interactions were examined by RNA pulldown, RNA immunoprecipitation, Tagged RNA affinity purification (TRAP) and electrophoretic mobility shift assays (EMSA).

**Results:** Here, we identified a functional rt-circRNA, termed rtcisE2F, that is highly expressed in liver cancer and liver TICs. rtcisE2F plays essential roles in the self-renewal and activities of liver TICs. rtcisE2F targets *E2F6* and *E2F3* mRNAs, attenuates mRNA turnover, and increases E2F6/E2F3 expression. Mechanistically, rtcisE2F functions as a scaffold of m^6^A reader IGF2BP2 and *E2F6/E2F3* mRNA, promotes the association of *E2F6*/*E2F3* mRNAs with IGF2BP2, and then inhibits their association with another m^6^A reader, YTHDF2. IGF2BP2 inhibits *E2F6*/*E2F3* mRNA decay, whereas YTHDF2 promotes *E2F6*/*E2F3* mRNA decay. By switching m^6^A readers, rtcisE2F enhances *E2F6*/*E2F3* mRNA stability. E2F6 and E2F3 are both required for liver TIC self-renewal and Wnt/β-catenin activation, and inhibition of these pathways is a potential strategy for preventing liver tumorigenesis and metastasis.

**Conclusion:** This work identified rtcisE2F as a key modulator in liver cancer and liver TICs, providing evidence for the biological function of rt-circRNA and unveiling a new regulatory layer for liver TIC self-renewal. rtcisE2F is involved in E2F6/E2F3 stability by switching their binding to the m^6^A readers IGF2BP2 and YTHDF2, providing a competitive mechanism between RNA molecules and m^6^A readers. Both E2F6 and E2F3 are required for liver TIC self-renewal and serve as therapeutic targets for liver TIC elimination.

## Introduction

Liver cancer is the third leading cause of cancer-related deaths with an increasing incidence, especially in China[1]. Despite of surgical resection, radiotherapy, chemotherapy and liver transplantation, many hepatocellular carcinoma (HCC) patients have a very poor prognosis, with high rates of relapse and metastasis. The failure of therapies is largely due to the significant heterogeneity of liver cancer[2]. Accumulating studies revealed that the broad heterogeneity is due to the hierarchic organization of tumor cells within the tumor that differentiate from a small subset of liver tumor initiating cells (TICs), also termed tumor initiating cells[2]. Several TIC markers have been identified, including CD133, CD13, CD44 and ALDH1. Unlike differentiated cancer cells, TICs harbor the ability to self-renew, differentiate and generate new tumors. Moreover, TICs are resistant to conventional therapies, including radiotherapy and chemotherapy drugs[3, 4]. Increasing studies demonstrate that TICs are also resistant to immunotherapy, including CAR-T and immune checkpoint therapy[5, 6]. Some novel technologies, including single-cell RNA sequencing and CRISPR-Cas9 based genome editing, have facilitated the characterization of TICs [7, 8]. However, the mechanisms involved in the self-renewal of liver TICs remain largely elusive.

Transcription factors (TFs) are sequence-specific DNA-binding factors and play central roles in cell fate determination. TFs bind to the promoter region of target genes to promote or inhibit their expression. E2F TFs are involved in cell cycle regulation and DNA synthesis in mammalian cells, and are also identified as modulators of cell fate in a number of stem cell lineages[9]. We previously identified ZIC2 as a functional TF in liver TICs[10]. However, the function and mechanism of TFs in liver TICs are still largely unknown.

N^6^-methyladenosine (m^6^A) is the most abundant modification of RNAs in humans[11]. The m^6^A modification is mediated by the METTL3/METTL14 methyltransferase complex, which comprises METTL3, METTL4, METTL14, WTAP and VIRMA[12], and removed by m^6^A demethylases, including FTO and AlKBH5[13, 14]. The function of RNA m^6^A modification is largely dependent on m^6^A readers, including YTHDF1, YTHDF2, YTHDF3, YTHDC1, YTHDC2, IGF2BP1, IGF2BP2, IGF2BP3, HNRNPA2B1 and HNRNPC[15]. m^6^A modification of RNAs is involved in many physiological and pathological processes, including spermatogenesis and embryonic development[16], DNA damage response[17], immunological regulation[18], and chromatin state[19]. In tumor cells, m^6^A modification contributes to tumorigenesis [20], metastasis [21] and drug resistance[22]. Recently, several m^6^A-related genes have been identified as modulators in various TICs, including bladder TICs[23], ovarian TICs[24] and glioblastoma stem cells[25].

Some stemness signaling pathways, including Wnt/β-catenin, Notch, Hedgehog and Hippo/Yap, are required for the self-renewal and maintenance of liver TICs, and the activation of these signaling pathways is precisely regulated[26, 27]. Regulatory RNAs have emerged as critical modulators in TIC self-renewal[28]. We previously reported that lnc β-catm and lncBRM regulate the self-renewal of liver TICs by modulating Wnt/β-catenin and Hippo/Yap1 signaling pathways[29, 30]. circRNAs, formed by covalent conjugation of 5’ and 3’ ends through backsplicing, emerges as critical modulators of various biological processes[31]. All circRNAs are generated from their parent pre-mRNAs, and grouped into extron circRNAs, intron circRNAs, and extron-intron circRNAs[32]. circRNAs play key roles in many biological processes, neuropsychiatric disorders, tumorigenesis and immunological responses[33, 34]. Several circRNAs, including ciRS-7/CDR1as, circHIPK3 and Sry circRNA, have been defined to act as miRNA sponges[35, 36]. Interestingly, fusion circRNAs, which cause cancer-associated chromosomal translocations, are involved in tumorigenesis and treatment resistance[37]. We also identified circPan3 and circKcnt2 regulate intestinal stem cell self-renewal and colitis progression[38, 39]. Recently, a novel class of circRNA, termed rt-circRNA, was identified as circRNA generated from the exons of two adjacent genes on the same strand[40]. However, the biological functions of rt-circRNA and the underlying molecular mechanisms are still unknown. Here we identified an rt-circRNA, named rtcisE2F (<u>rt</u>-<u>ci</u>rcRNA <u>s</u>tabilizing *<u>E2F</u>6*/*E2F3* mRNAs), that was highly expressed in liver cancer and liver TICs. rtcisE2F modulated the association of *E2F6*/*E2F3* mRNAs with the m^6^A readers IGF2BP2 and YTHDF2, which promote and inhibit *E2F6*/*E2F3* mRNA stability, respectively. Our work reveals a new regulatory mechanism of circRNA via m^6^A modification, and also shows E2F6/E2F3 as a target for eliminating liver TICs.

## Results

### rtcisE2F is highly expressed in liver cancer and liver TICs

rt-circRNAs are a novel class of circRNAs that are derived from exons originating from two adjacent genes on the same strand [40]. However, the biological function and molecular mechanism of rt-circRNA are unknown. To identify functional rt-circRNA in liver tumorigenesis and liver TICs, we analyzed the rt-circRNA expression pattern in liver cancer, and screened out eight rt-circRNAs expressed in liver cancer (Supplementary Fig. 1A). Among these eight rt-circRNAs, four transcripts were highly expressed in CD133^+^ liver TICs (Supplementary Fig. 1B). These four rt-circRNAs were further validated by RNase R digestion and actinomycin D treatment (Supplementary Fig. 1C). rt-circRNA knockdown cells were established, and attenuated sphere formation capacity was detected only in rtcisE2F-silenced cells (Figure 1A and Supplementary Fig. 1D). rtcisE2F was highly expressed in liver cancer, particularly in advanced liver cancer (Figure 1B-D), and its expression was correlated with the relapse and prognosis of HCC (Figure 1E, F). rtcisE2F expression levels were positively correlated with the CD133^+^ TIC ratio and CD133 expression, indicating the high expression of rtcisE2F in liver TICs (Figure 1G, H). Indeed, rtcisE2F was highly expressed in CD133^+^ liver TICs and spheres (Figure 1I, J). Interestingly, rtcisE2F expression was negatively correlated with the expression of its parent genes (*CYP2C18* and *CYP2C19*) (Supplementary Fig. 1E). Overall, these findings demonstrate that rtcisE2F, an rt-circRNA, is highly expressed in liver cancer and liver TICs.

**Figure 1.**
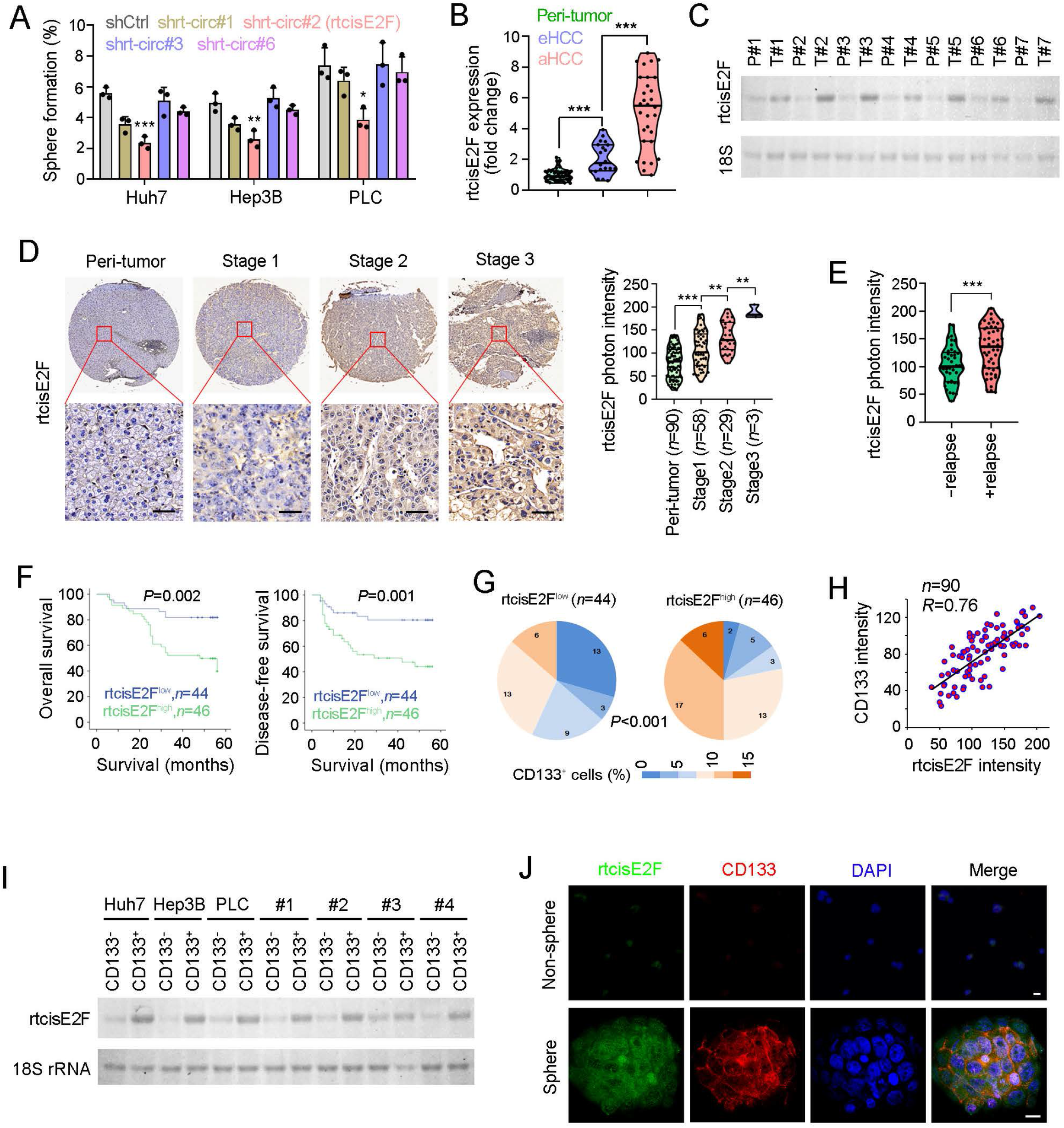
High expression of rtcisE2F in liver cancer and TICs. (A) Sphere formation of 1000 rt-circRNA silenced and control cells. rt-circ#2 denotes rtcisE2F. *n*=3 independent experiments. (B) Quantitative real-time PCR analysis for rtcisE2F expression in 50 peri-tumor, 20 early hepatocellular carcinoma (eHCC) and 30 advanced hepatocellular carcinoma (aHCC). All expression levels were normalized to the average level of rtcisE2F in peri-tumor samples. (C) Northern blot of rtcisE2F in paired liver tumors (T) and peri-tumors (P). *n*=7 samples were examined and 18S rRNA was a loading control. (D) rtcisE2F *in situ* hybridization of liver cancer tissue assay containing 90 peri-tumor, 58 stage 1, 29 stage 2, and 3 stage 3 tumor tissues. Left panels are representative images and right panels are violin plot, in which medium, minimum, maximum and quarter levels were shown, as well as individual samples. Scale bars, 50 μm. (E) Violin plot of rtcisE2F expression in HCC samples with relapse (+relapse) or without relapse (-relapse). (F) Kaplan–Meier survival analysis of rtcisE2F highly expressed (green) and lowly expressed (blue) samples, which were grouped according to average levels of rtcisE2F intensity in 90 liver cancer tissues. (G) Percentage distribution of CD133^+^ TICs in rtcisE2F^low^ (left) and rtcisE2F^high^ (right) samples. (H) Co-expression of rtcisE2F and liver TIC marker CD133 in 90 liver cancer tissues. (I) Northern blot of rtcisE2F in CD133^+^ liver TICs and CD133^−^ liver non-TICs, which were enriched rom HCC cell lines or primary samples via FACS. 18 rRNA is a loading control. (J) Fluorescence *in situ* hybridization (FISH) of rtcisE2F in spheres and non-spheres. Scale bars, 10 μm. In all panels, data are shown as mean ± s.d. \**P* < 0.05; **, *P*<0.01; \*\*\**P* < 0.001, by one-tailed Student’s T-test.

### rtcisE2F drives self-renewal of liver TIC

To elucidate the role of rtcisE2F in liver TICs, we deleted rtcisE2F in HCC cells. circRNA formation depends on intronic complementary sequences flanking exons, providing a possible strategy for circRNA knockout [32, 39]. Through a mini-gene analysis, we generated *rtcisE2F* knockout cells using CRISPR-Cas9, and confirmed the deletion of rtcisE2F (Figure 2A and Supplementary Fig. A-D). *rtcisE2F* knockout cells showed impaired propagation, sphere formation and metastatic activities, confirming the critical role of rtcisE2F in liver TIC self-renewal (Figure 2B-D). Moreover, *rtcisE2F* knockout cells showed attenuated tumor initiating capacity, confirming the essential role of rtcisE2F in the function of liver TICs (Figure 2E, F).

**Figure 2.**
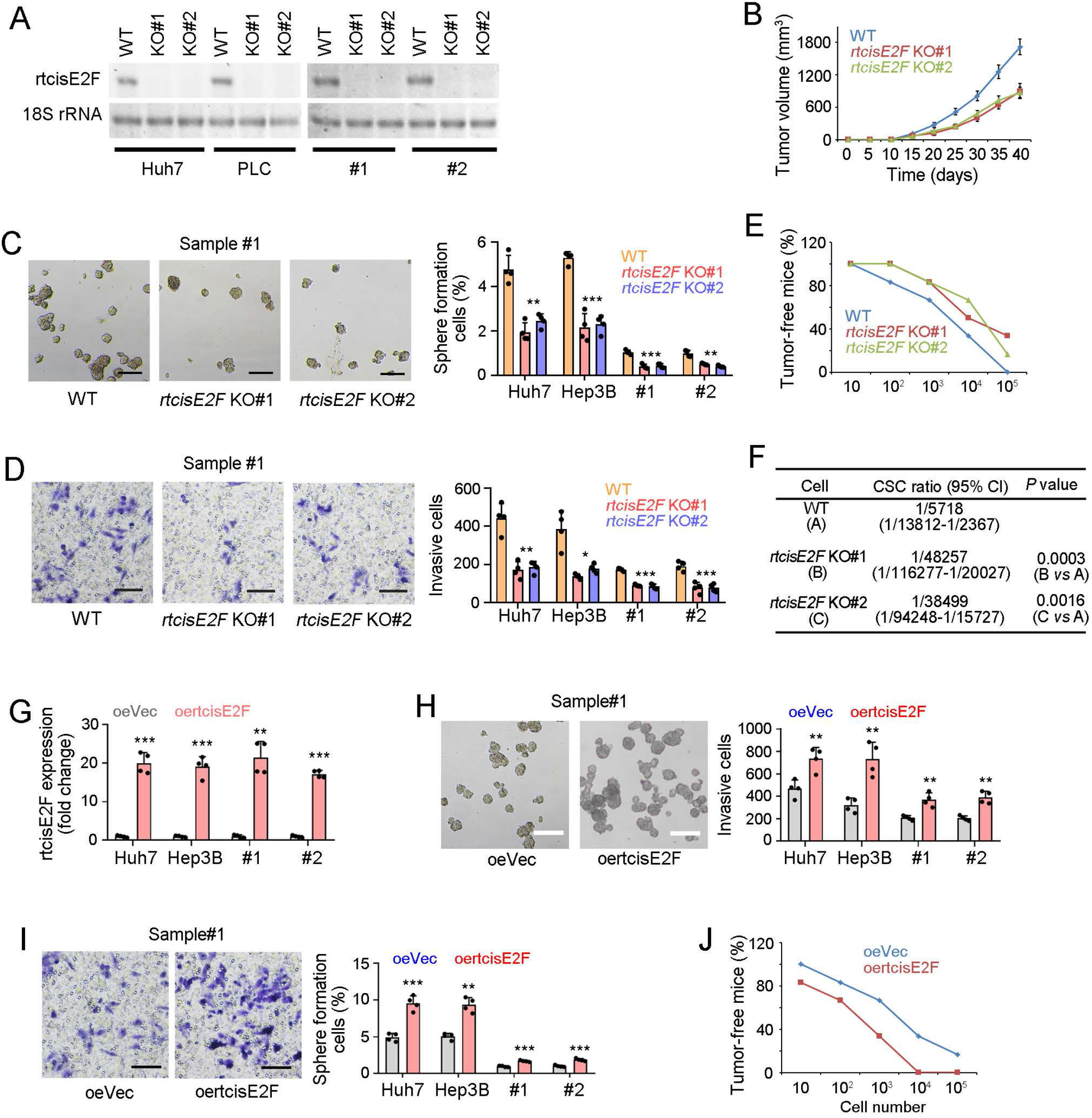
rtcisE2F is required for liver TIC self-renewal. (A) Northern blot to validate *rtcisE2F* KO efficiency in *rtcisE2F* knockout cells generated via CRISPR-Cas9 approach. #1, primary sample #1; #2, primary sample #2. (B) Tumor propagation of *rtcisE2F* knockout cells. The indicated cells were subcutaneously injected into BALB/c nude mice and tumor volume was detected at indicated days after injection. (C) Sphere formation assay of *rtcisE2F* knockout cells. *n*=1000 Huh7, Hep3B cells, and *n*=5000 primary cells were used for sphere formation. Scale bars, 500 μm. (D) Transwell assay of *rtcisE2F* knockout cells, showing representative images (left) and invasive ratios (right). Scale bars, 70 μm. (E, F) Tumor initiation assay of *rtcisE2F* knockout cells using gradient numbers of indicated cells. *n*=6 mice for each group and tumor formation was monitored 3 months after transplantation (E). TIC ratios calculated by extreme limiting dilution analysis (http://bioinf.wehi.edu.au/software/elda/) were in (F). 95% CI, 95% confidence interval of the estimation; vs, versus. (G) Quantitative real-time PCR confirmation for rtcisE2F overexpression. oeVec, overexpressing empty vector; oertcisE2F, overexpressing rtcisE2F. (H, I) Sphere formation (H) and transwell assay (I) of rtcisE2F overexpressing cells. Scale bars, H, 500 μm; I, 70 μm. (J) Tumor initiation assay of gradient numbers of oeVec and oertcisE2F cells. *n*=6 mice were subcutaneously injected with gradient numbers of indicated cells for 3 months’ tumor formation. In all panels, data are shown as mean ± s.d. \**P* < 0.05; \*\**P* < 0.01; \*\*\**P* < 0.001, by one-tailed Student’s T-test. At least three independent experiments were performed with similar results.

We also generated rtcisE2F overexpressing cells, which showed enhanced sphere formation and metastatic activities, confirming the key role of rtcisE2F in the self-renewal of liver TICs (Figure 2G-I). Tumor initiation was also enhanced in rtcisE2F overexpressing cells (Figure 2J). Taken together, these findings confirm that rtcisE2F plays critical roles in the self-renewal and maintenance of liver TICs.

### rtcisE2F targets E2F6 and E2F3 to drive liver TIC self-renewal

To explore the molecular mechanism of rtcisE2F in liver TIC self-renewal, we analyzed its potential target genes. Considering the critical role of TFs in cell fate determination, we analyzed the coexpression of TFs and rtcisE2F parent genes (Supplementary Fig. 3A). Among the top 20 coexpressed TFs, the expression levels of eight TFs were significantly altered in *rtcisE2F* knockout cells (Supplementary Fig. 3B). Therefore, we depleted these eight TFs through CRISPR interference (CRISPRi) (Supplementary Fig. 3C). Two of these TFs, E2F6 and E2F3, were required for sphere formation, suggesting that E2F6 and E2F3 are functional targets for rtcisE2F (Figure 3A). In fact, the expression levels of E2F6 and E2F3 were reduced in *rtcisE2F* knockout cells (Figure 3B). The coexpression of rtcisE2F and E2F6/E2F3 was also confirmed in HCC samples (Figure 3C). Overall, these data demonstrate that E2F6 and E2F3 are target genes for rtcisE2F in liver TICs.

**Figure 3.**
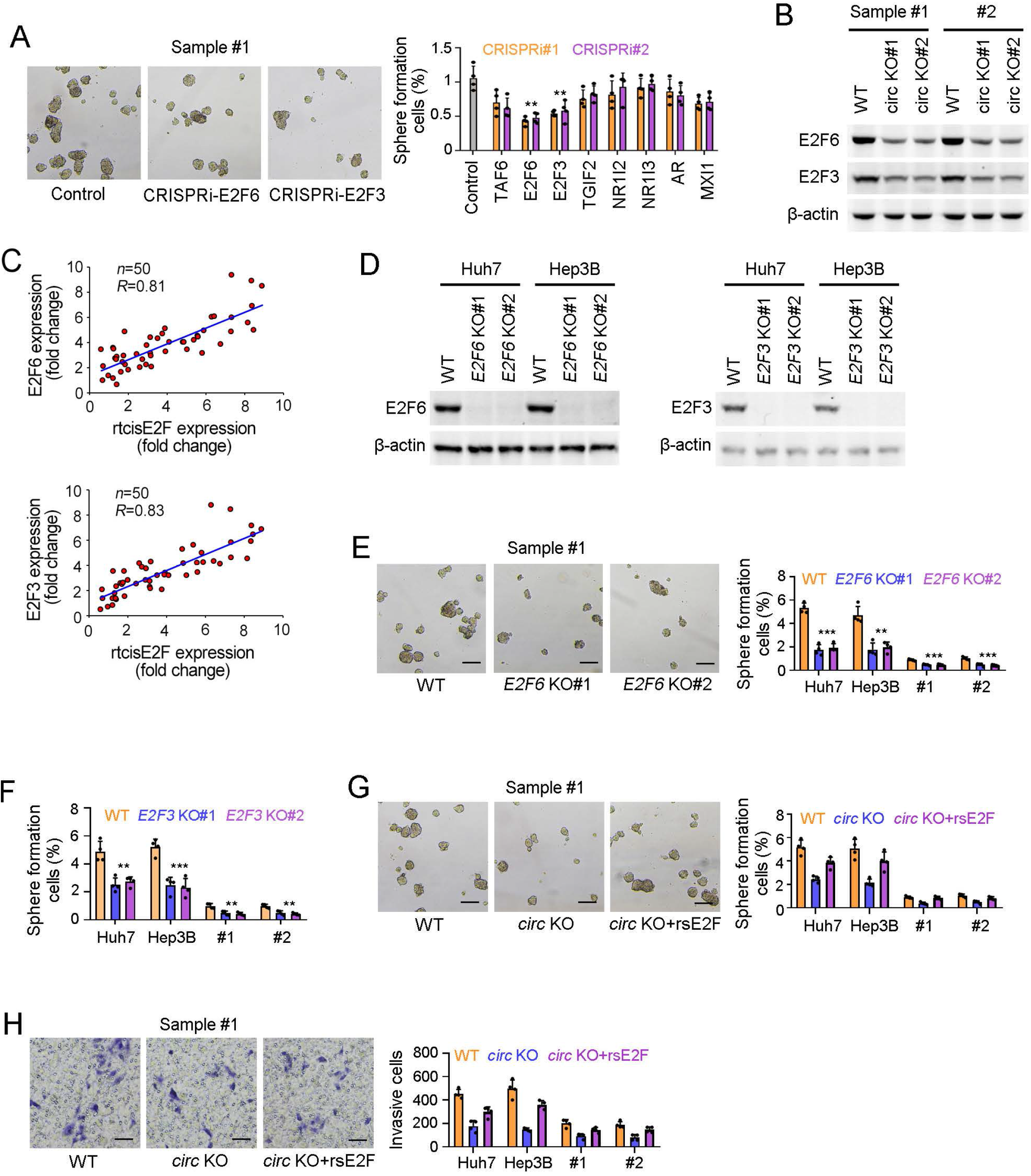
rtcisE2F targets E2F6 and E2F3 to drive liver TIC self-renewal. (A) Sphere formation of indicated TF depleted cells, which were generated via CRISPRi-KRAB strategy. (B) Western blot for E2F6 and E2F3 expression in *rtcisE2F* knockout cells. β-actin is a loading control. (C) Co-expression of rtcisE2F with E2F6 (upper) or E2F3 (lower). 50 samples were detected and expression levels were normalized to the average levels in peri-tumors. (D) Western blot to validate *E2F6* and *E2F3* knockout using indicated KO and control cells, which were generated via CRISPR-Cas9 approach. (E, F) Sphere formation assay of *E2F6* (E) and *E2F3* (F) knockout cells. *n*=1000 cells (for Huh7 and Hep3B) and *n*=5000 cells (for primary cells) were used for sphere formation. (G, H) Sphere formation (G) and transwell assay (H) of E2F rescued *rtcisE2F* knockout cells. E2F6 and E2F3 were both rescued in *rtcisE2F* knockout cells, which were named circ KO+rsE2F cells. In all panels, data are shown as mean ± s.d. \*\**P* < 0.01; \*\*\**P* < 0.001, by one-tailed Student’s T-test. For all representative images, at least three independent experiments were performed with similar results.

We next generated *E2F6* knockout and *E2F3* knockout cells using CRISPR-Cas9, and western blotting confirmed the knockout was successful (Figure 3D). *E2F6* and *E2F3* knockout cells showed impaired sphere formation capacities, indicating their essential role in liver TIC self-renewal (Figure 3E, F). Rescuing E2F6 and E2F3 expression in *rtcisE2F* knockout cells restored the sphere formation and metastasis capacities, demonstrating that the function of rtcisE2F in liver TICs is mediated by E2F6 and E2F3 (Figure 3G, H). Thus, rtcisE2F drives the self-renewal of liver TICs via the TFs E2F6 and E2F3.

### rtcisE2F combines with *E2F6*/*E2F3* mRNAs to maintain their stability

To further explore the molecular mechanism of rtcisE2F in E2F6 and E2F3 expression, we first performed ChIRP assay, but rtcisE2F did not bind to *E2F6*/*E2F3* promoter (Figure 4A and Supplementary Fig. 4A). Meanwhile, there were no difference of transcription activity and *E2F6* promoter activity between *rtcisE2F* knockout and control cells (Figure 4B, C). These results indicate that rtcisE2F is not involved in E2F6/E2F3 transcription.

**Figure 4.**
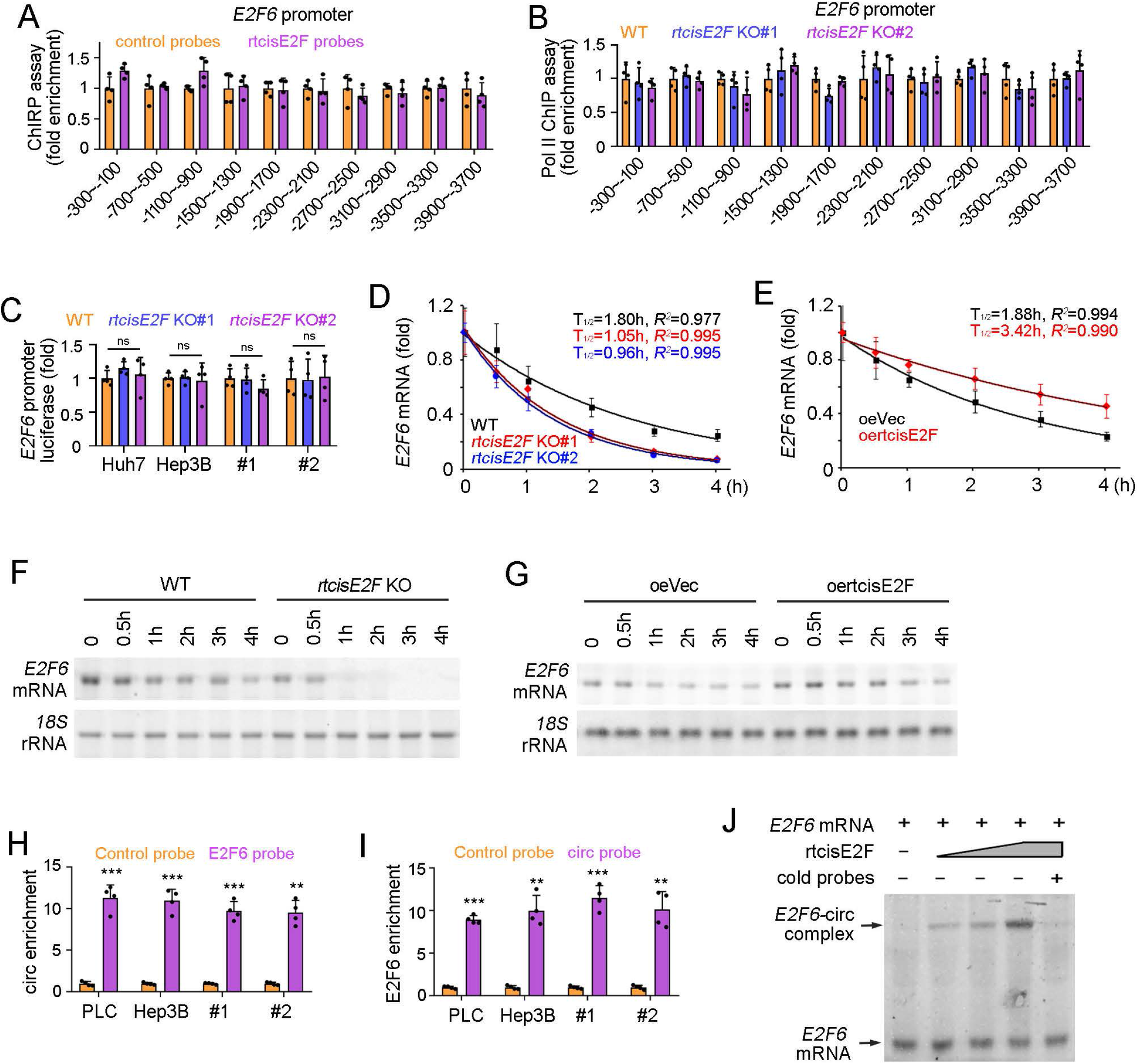
rtcisE2F promotes *E2F6*/*E2F3* mRNA stability. (A) Quantitative real-time PCR for enrichment of *E2F6* promoter in eluate from Chromatin Isolation by RNA purification (ChIRP) with rtcisE2F probes. (B) Chromatin immunoprecipitation (ChIP) assay of RNA polymerase II (Pol II) from *rtcisE2F* knockout cells, followed by quantitative real-time PCR analysis for the enrichment of *E2F6* promoter. (C) Luciferase activity of *E2F6* promoter (−4000bp~0bp) in *rtcisE2F* knockout cells, which were normalized to luciferase activity in WT cells. (D, E) Quantitative real-time PCR analysis of *E2F6* mRNA in *rtcisE2F* knockout (KO) TICs (D) and rtcisE2F overexpressing (oe) TICs (E), which were treated with 2 μg/ml actinomycin D for the indicated times. T_1/2_, half time. (F, G) Northern blot of *E2F6* mRNA in *rtcisE2F* knockout TICs (F) and rtcisE2F overexpressing TICs (G) treated with 2 μg/ml actinomycin D for indicated time points. (H, I) Quantitative real-time PCR analysis of RNA pulldown eluate to detect the interaction of rtcisE2F and *E2F6* mRNA. (J) RNA mobility-shift assay of gradient rtcisE2F with *E2F6* mRNA. In all panels, data are shown as mean ± s.d. \*\**P* < 0.01; \*\*\**P* < 0.001; ns, not significant, by one-tailed Student’s T-test. For all representative images, at least three independent experiments were performed with similar results.

Considering that rtcisE2F regulates *E2F6*/*E2F3* mRNA levels but not their transcription, we determined the effect of rtcisE2F on *E2F6*/*E2F3* mRNA stability. As expected, impaired *E2F6* and *E2F3* mRNA stability was observed in *rtcisE2F* knockout cells (Figure 4D, and Supplementary Fig. 4B). On the contrary, rtcisE2F overexpressing cells showed enhanced *E2F6* and *E2F3* mRNA stability (Figure 4E, and Supplementary Fig. 4C). The involvement of rtcisE2F in *E2F6* mRNA stability was confirmed by Northern blot (Figure 4F, G). Then the interaction between rtcisE2F and *E2F6*/*E2F3* mRNAs was detected using RNA pulldown and confirmed by EMSA (Figure 4H-J, and Supplementary Fig. 4D, E). Overall, these findings indicate that rtcisE2F binds to *E2F6*/*E2F3* mRNAs and inhibits their turnover, maintaining their stability.

### rtcisE2F switches the binding of m^6^A readers IGF2BP2 and YTHDF2 to *E2F6*/*E2F3* mRNAs

We further investigated the molecular mechanism by which rtcisE2F maintains *E2F6*/*E2F3* mRNA stability by performing TRAP assay[41] (Supplementary Fig. 5A). In *rtcisE2F* knockout spheres, we found decreased binding of *E2F6* mRNA to IGF2BP2 but increased binding to YTHDF2 according to mass spectrum, with confirmation by Western blot (Figure 5A, B). RIP assay also demonstrated the impaired IGF2BP2-E2F6/E2F3 binding and enhanced YTHDF2-E2F6/E2F3 binding in *rtcisE2F* knockout cells, and addition of gradient rtcisE2F elicited opposite effects (Figure 5C-E, and Supplementary Fig. 5B, C). These data demonstrated that rtcisE2F is involved in the interactions of IGF2BP2/YTHDF2 with *E2F6*/*E2F3* mRNAs.

**Figure 5.**
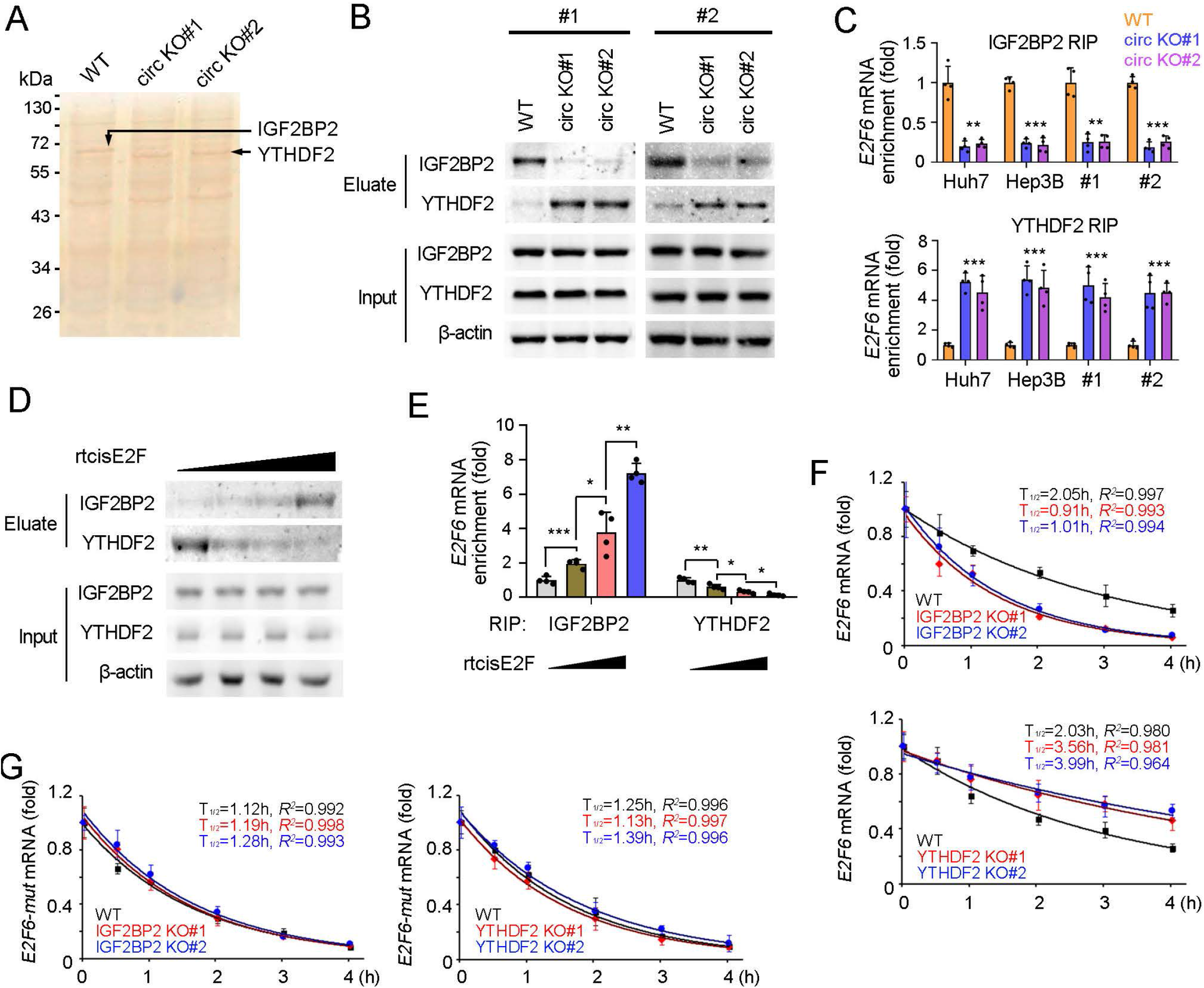
rtcisE2F participates in the interaction of *E2F6*/*E2F3* mRNAs with IGF2BP2/YTHDF2. (A) Silver staining of the *E2F6* mRNA pulldown eluate from *rtcisE2F* KO and control spheres, which were derived from primary HCC samples. IGF2BP2 and YTHDF2 were identified via Mass Spectrum. (B) Western blot of IGF2BP2 and YTHDF2 in the eluate of TRAP assay, which was performed using *rtcisE2F* knockout and control spheres. (C) Quantitative real-time PCR of *E2F6* mRNA in RNA immunoprecipitation (IP) eluate using IGF2BP2 antibody (upper panel) and YTHDF2 antibody (lower panel). *rtcisE2F* knockout and WT spheres were used for RIP, and *E2F6* mRNA enrichment levels were normalized to those in WT cells. (D) Western blot of IGF2BP2 and YTHDF2 in RNA pulldown eluate using E2F6 probes and sphere lysate, supplemented with gradient rtcisE2F transcripts. (E) Quantitative real-time PCR of *E2F6* mRNA in RNA immunoprecipitation (RIP) eluate using IGF2BP2/YTHDF2 antibodies and sphere lysate supplemented with gradient rtcisE2F transcripts. (F) Quantitative real-time PCR analysis of *E2F6* mRNA in *IGF2BP2* knockout TICs (up) and *YTHDF2* knockout TICs (down), which were treated with 2 μg/ml actinomycin D for the indicated times. (G) Stability of mutant *E2F6* mRNA, which harbors no m^6^A site, was evaluated via quantitative real-time PCR as F. In all panels, data are shown as mean ± s.d. \**P* < 0.05; \*\**P* < 0.01; \*\*\**P* < 0.001, by one-tailed Student’s T-test. For all representative images, at least three independent experiments were performed with similar results.

IGF2BP2 and YTHDF2 are well-known m^6^A readers, which typically inhibits and promotes the decay of target RNA molecules, respectively[42, 43]. We generated *IGF2BP2* and *YTHDF2* knockout cells using CRISPR-Cas9 (Supplementary Fig. 5D). *E2F6*/*E2F3* mRNA stability was impaired in *IGF2BP2* knockout cells, but enhanced in *YTHDF2* knockout cells (Figure 5F, and Supplementary Fig. 5E). We also generated mutant *E2F6* mRNA lacking m^6^A modification site, and revealed that the stability of this mRNA was unaffected by knockout of IGF2BP2 or YTHDF2 (Figure 5G, and Supplementary Fig. 5F, G). Taken together, these findings indicate that rtcisE2F switches IGF2BP2/YTHDF2 binding to the m^6^A site in E2F6/E2F3 to maintain the stability of *E2F6*/*E2F3* mRNAs.

### rtcisE2F promotes the interaction of IGF2BP2 and E2F6/E2F3

We then investigated the molecular mechanism of rtcisE2F in binding switches of IGF2BP2 and YTHDF2 to E2F6/E2F3. We first analyzed their interaction, and found that rtcisE2F interacted with IGF2BP2, but not YTHDF2 (Figure 6A, B). Through domain mapping and TRAP assay, we identified that second exon (originating from exon 9 of the CYP2C18) was necessary to bind IGF2BP2 (Figure 6C, and Supplementary Fig. 6A). We next predicted the tertiary structure of second exon, and identified that hairpin region 9 (HR9) was necessary to associate with IGF2BP2 (Figure 6D, Supplementary Fig. 6B). Through TRAP assay and real-time PCR analysis, we identified the HR5 in the third exon (originating from exon 1 of the CYP2C19) was necessary for rtcisE2F binding to *E2F6/E2F3* (Figure 6E, F, and Supplementary Fig. 6C). We also constructed mutant rtcisE2F, which can’t bind to IGF2BP2 or *E2F6/E2F3* because of HR9 or HR5 mutation (Supplementary Fig. 6B, C). Both mutant rtcisE2F transcripts were not involved in *E2F6/E2F3* mRNA regulation, indicating that rtcisE2F exerts its role through interaction with IGF2BP2 and *E2F6/E2F3* mRNA (Figure 6G). Of note, nucleotide fragment of *E2F6/E2F3* mRNA is almost totally paired with HR5 of rtcisE2F (Supplementary Fig. 6D). We also constructed *E2F6/E2F3* mutations that can’t combine with recisE2F, and demonstrated that rtcisE2F had no role in mRNA stability of mutant *E2F6/E2F3* (Figure 6H and Supplementary Fig. 6E). Altogether, rtcisE2F promotes the interaction of IGF2BP2 and *E2F6/E2F3* mRNA as a scaffold, and finally drives the binding switch of IGF2BP2 and YTHDF2.

**Figure 6.**
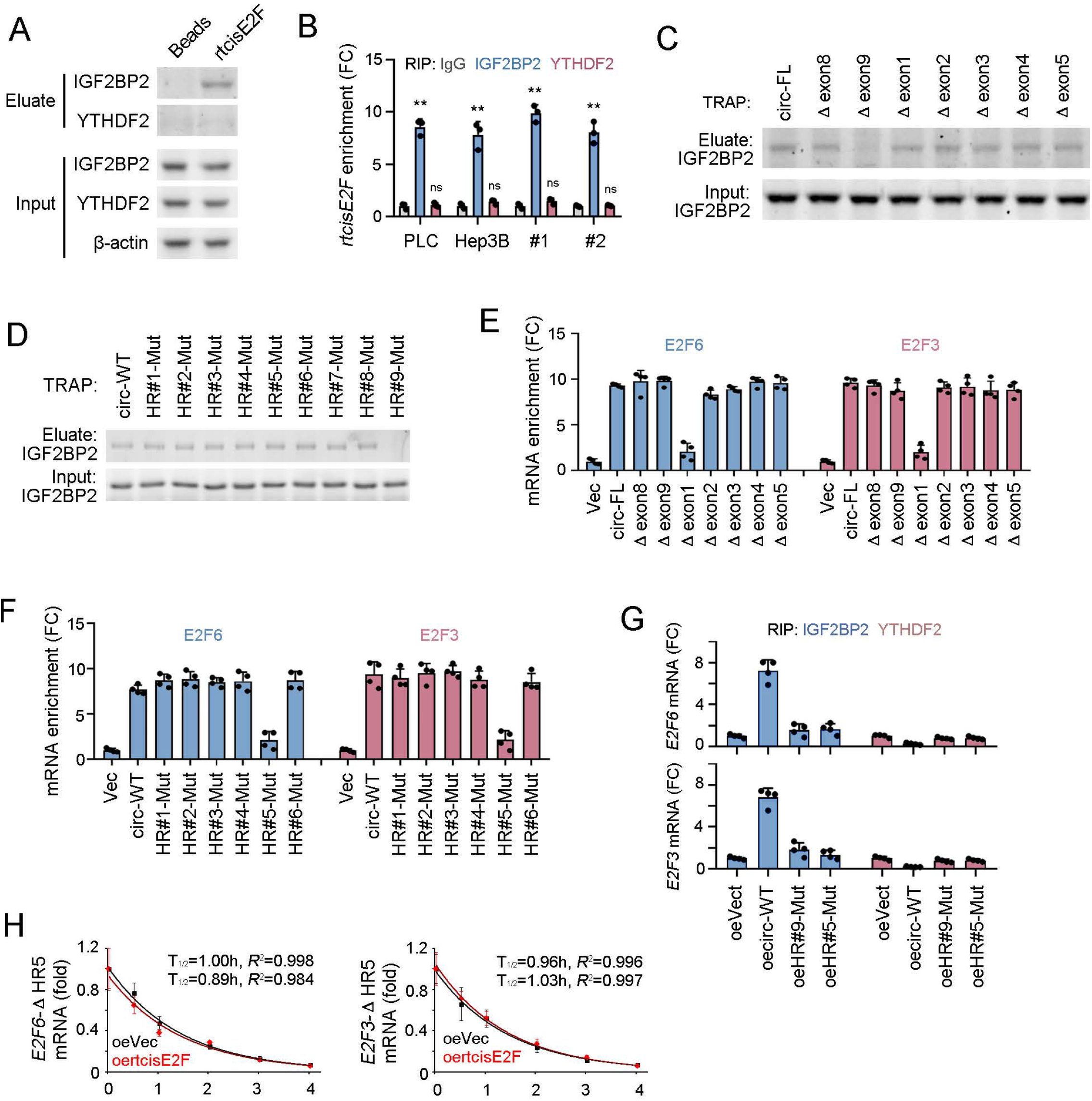
rtcisE2F functions as a scaffold to promote the interaction of IGF2BP2 and *E2F6/E2F3* mRNAs. (A) Western blot for IGF2BP2 and YTHDF2 in RNA pulldown eluate, in which sphere lysate was used. (B) Quantitative real-time PCR analysis for rtcisE2F enrichment in RNA immunoprecipitation (RIP) eluate. Anti-IGF2BP2, anti-YTHDF2 and control antibodies were used for RIP. FC, fold change. (C, D) Western blot for IGF2BP2 enrichment in TRAP eluate, in which MS2 conjugated truncated rtcisE2F (C) or MS2 conjugated mutant rtcisE2F (D) were used. (E, F) Quantitative real-time PCR for *E2F6* and *E2F3* mRNA enrichment in the eluate of TRAP assay. All enrichment levels were normalized to those in control cells. (G) Quantitative real-time PCR for *E2F6* and *E2F3* mRNA levels in the indicated cells, which overexpress wild-type (WT) rtcisE2F and mutant rtcisE2F. All expression levels were normalized to those in control cells. (H) Quantitative real-time PCR to detect the stability of *E2F6*-ΔHR5 and *E2F3*-ΔHR5 in rscisE2F overexpressing and control cells. Δ HR5, losing HR5 interaction ability. For all representative images, at least three independent experiments were performed with similar results.

### E2F6 and E2F3 are involved in liver tumorigenesis and may serve as therapeutic targets

As shown above, E2F6 and E2F3 are required for liver TIC self-renewal. We further explored whether E2F6 and E2F3 are involved in liver tumorigenesis and severity. We first analyzed E2F6/E2F3 expression levels using Wang’s cohort (GSE14520[44, 45]), and confirmed that E2F6 and E2F3 are highly expressed (they are the top two E2F TFs expressed) in liver cancers (Supplementary Fig. 7A, B). High expression of E2F6 and E2F3 in primary liver cancer samples was also confirmed by quantitative real-time PCR and Western blot (Figure 7A, B). Moreover, there was higher expression of E2F6 and E2F3 in spheres than in non-spheres (Figure 7C). The expression levels of E2F6 and E2F3 were also related to the clinical severity, metastasis and prognosis of liver cancer (Figure 7D and Supplementary Fig. 7C, D). These data demonstrate that E2F6 and E2F3 are highly expressed in liver cancer and liver TICs.

**Figure 7.**
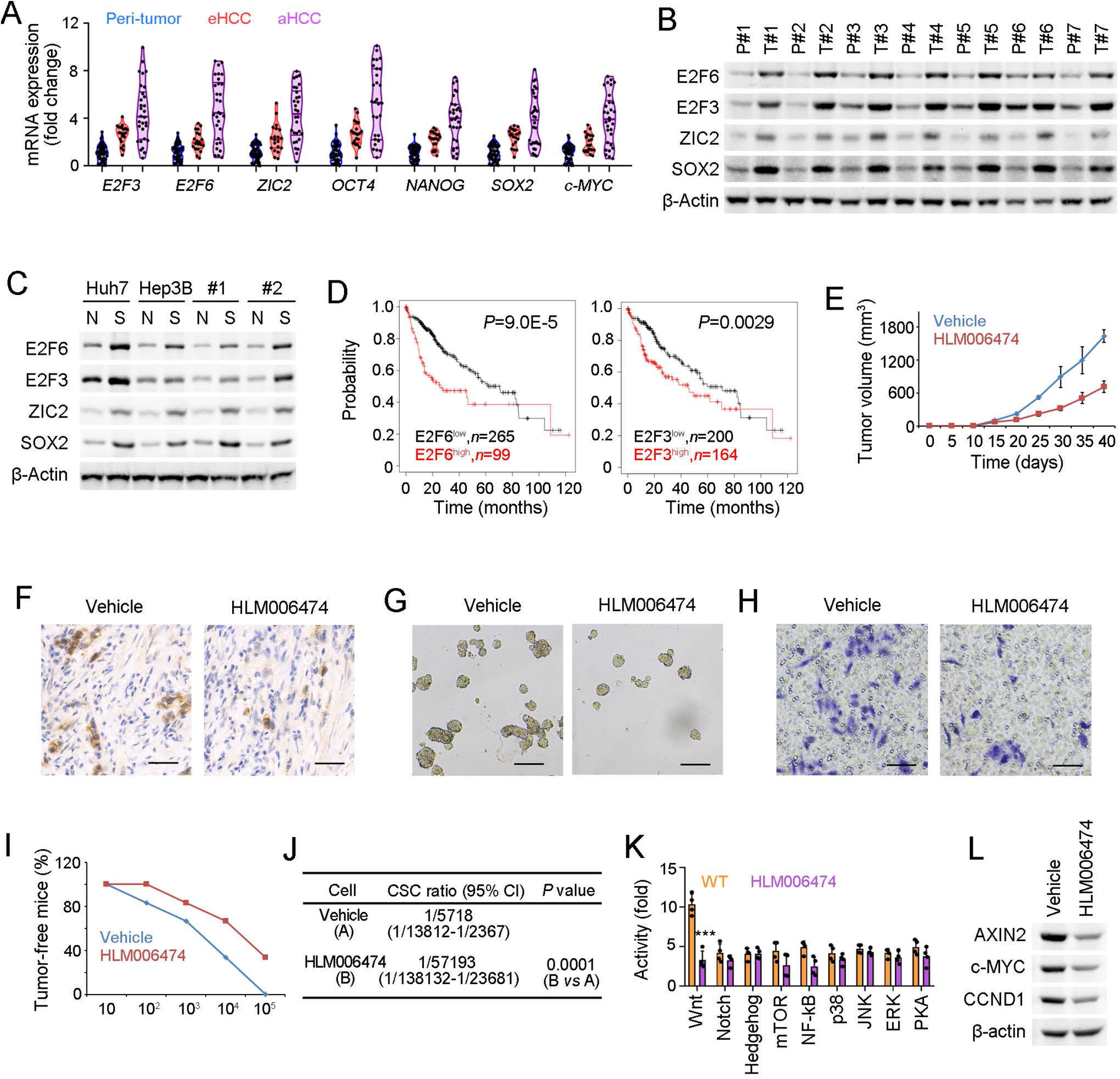
E2F6 and E2F3 serve as therapeutic targets for liver TIC elimination. (A) Violin plots showing mRNA expression levels of *E2F3*, *E2F6* and several TIC-related TFs, in which individual expressions are shown as black dots, with medium, minimum, maximum and quarter levels shown as lines. 50 peri-tumors, 20 early HCC (eHCC) and 30 advanced HCC (aHCC) samples were detected. (B) Western blot for indicated TF expression in liver tumor (T) and peri-tumor (P) samples. (C) Western blot to detect TF expression in non-sphere (N) and sphere (S) samples, which were derived from Huh7, Hep3B and primary samples. (D) Kaplan–Meier survival analysis of E2F6 and E2F3 highly expressed and lowly expressed samples using online-available data (http://kmplot.com/analysis/). (E) Tumor propagation of primary liver cancer cells treated with E2F inhibitor HLM006474, which were subcutaneously injected into BALB/c nude mice. (F) CD133 immunohistochemistry in HLM006474 treated and control xenografts. Scale bars, 100 μm. (G, H) Sphere formation (G) and transwell assay (H) of tumor cells isolated from HLM006474 treated and control xenografts. (I, J) 3 months’ tumor initiation assay using gradient numbers of liver cancer cells, which were from HLM006474 treated and control xenografts. *n*=6 mice for each group and the ratios of tumor-free mice were shown in (I) and TIC ratios calculated by extreme limiting dilution analysis (J). 95% CI, 95% confidence interval of the estimation; vs, versus. (K) FACS detection for the activity of indicated signaling pathways in HLM006474 treated and control primary cells. (L) Western blot for Wnt/β-catenin activation in HLM006474 treated and control cells. In all panels, data are shown as mean ± s.d. \**P* < 0.05; \*\**P* < 0.01; \*\*\**P* < 0.001, by one-tailed Student’s T-test. For all representative images, at least three independent experiments were performed with similar results.

We then investigated the therapeutic potential of E2F6 and E2F3 for liver cancer. Tumor propagation was impaired in liver cells treated with HLM006474, a pan-E2F TF inhibitor [46] (Figure 7E). Interestingly, the HLM006474-treated tumors contained fewer CD133^+^ TICs and displayed impaired sphere formation and transwell capacities (Figure 7F-H). Moreover, HLM006474 treatment impaired tumor initiation, confirming the therapeutic role of inhibiting E2F in eliminating liver TICs (Figure 7I, J). Finally, we evaluated the signaling pathways involved in the effects of HLM006474, and found that Wnt/β-catenin signaling, one of the most important signaling pathways in the self-renewal of liver TICs[47], was significantly impaired in HLM006474 treated cells (Figure 7K). The role of HLM006474 in targeting the Wnt signaling pathway was confirmed by western blotting (Figure 7L). Taken together, these findings demonstrate that E2F6/E2F3 drives liver tumorigenesis and can serve as a therapeutic target for liver cancer and the elimination of liver TICs.

## Discussion

In this work, we revealed that the rtcisE2F-IGF2BP2/YTHDF2-E2F6/E2F3 signaling pathway is activated in liver TICs. rtcisE2F promotes IFG2BP2 binding and inhibits YTHDF2 binding to m^6^A-modified *E2F6*/*E2F3* mRNAs, inhibits *E2F6*/*E2F3* mRNA turnover and finally drives the self-renewal of liver TICs (Supplementary Fig. 7E). Moreover, E2F6 and E2F3 are therapeutic targets for liver tumorigenesis and liver TIC elimination. Our work reveals a previously unknown molecular mechanism driving self-renewal of liver TICs, and adds a new layer of circRNA function in target mRNA recognition by m^6^A readers.

circRNAs originate from their parent RNAs via covalent conjugation of the 5’ and 3’ ends, thus circRNAs and their corresponding linear mRNAs are derived from the same parent transcripts. However, the relationship between circRNA expression and the corresponding linear RNA expression is unclear[48]. In some studies, there was no correlation between circRNA and linear RNA expression[39], but positive or negative correlations were detected in some other studies. For example, cyclization of circACC1, a circRNA associated with colorectal cancer metabolism, inhibits splicing of linear ACC1 RNA[49]. Here, we showed that rtcisE2F expression was negatively correlated with the expression levels of the linear genes CYP2C18 and CYP2C19, although the molecular mechanisms involved in this negative correlation require further investigation.

There are many different subclasses of circRNA and accumulating circRNA types have been identified on account of the advances in RNA extraction, library construction, RNA sequencing and analysis. For example, nuclear EIciRNAs (exon-intron type circRNAs) are involved in gene transcription[50], while mitochondrial mecciRNAs regulate the mitochondrial location of associated proteins[51]. rt-circRNA is a newly identified type of circRNA that is formed from exons originating from two adjacent genes on the same strand[40]. Although rt-circRNAs are expressed in some tumors, their roles in tumorigenesis, metastasis and TIC self-renewal have not been studied, and their regulatory mechanisms are unknown. Here, we identified rtcisE2F, a functional rt-circRNA that regulates the self-renewal and metastatic capacities of liver TICs. circRNAs exert their roles via various mechanisms, involving gene transcription, protein stability, subcellular localization, protein activity, alternative splicing and so on[52]. In liver TICs, rtcisE2F directly binds to *E2F6*/*E2F3* mRNAs, participates in the assembly of mRNAs and RNA binding proteins (RBP) IGF2BP2 and YTHDF2, and ultimately inhibits the degradation of *E2F6* and *E2F3* mRNAs. In fact, we previously revealed that lncBRM is involved in the assembly of a protein complex. lncBRM inhibits the assembly of the BRG1-type SWI/SNF complex but promotes the assembly of the BRM-type complex, leading to the self-renewal of liver TICs[30]. In this study, we identified that rt-circRNA, a new kind of circRNA, is involved in mRNA-RBP assembly and liver TIC self-renewal, adding a new layer to circRNA function and their regulatory mechanisms.

As the most abundant RNA modification in mammalian mRNA, m^6^A plays a key role in many physiological and pathological processes[53]. RNA can be methylated by m^6^A writers, demethylated by m^6^A erasers, and recognized by m^6^A readers. The m^6^A modification is involved in the stability and translation of target RNAs[42, 54]. The functionally distinct effects of m^6^A modification are partially mediated by the different readers that bind to m^6^A RNA molecules. Classically, YTHDF1 binds m^6^A mRNAs to regulate protein translation activity[54], whereas YTHDF2 binds m^6^A RNA molecules to promote RNA degradation[42]. More recently, it has been recognized that the IGF2BP family of RBPs act as m^6^A readers that bind to m^6^A RNA molecules and inhibit mRNA degradation[43]. Considering the opposite effects of YTHDF2 and IGF2BPs on RNA stability, it is important to determine which of these readers binds to m^6^A mRNA. However, the m^6^A RNA-m^6^A reader interaction and the competition between various m^6^A readers have not been well studied. Here, we showed that rtcisE2F is involved in the interaction between m^6^A mRNA and m^6^A readers, by promoting the binding of E2F6/E2F3 RNA to IGF2BP and inhibiting their binding to YTHDF2, and finally enhancing the stability of *E2F6*/*E2F3* mRNAs. This research has uncovered the regulatory mechanism of m^6^A mRNA stability, and there is still much work to be done, such as the molecular mechanism by which rtcisE2F regulates the reorganization of *E2F6*/*E2F3* mRNAs by the m^6^A readers YTHDF2/IGF2BP2, and the combinations of m^6^A readers with other m^6^A RNA molecules.

TFs play a central role in cellular fate determination. Like normal stem cells, TICs are regulated by well-known stemness TFs, such as c-MYC, OCT4, NANOG, and SOX2[10]. We previously revealed that the TF ZIC2 is highly expressed in liver cancer and liver TICs, recruits the NURF chromatin remodeling complex to the *OCT4* promoter, and drives the self-renewal of liver TICs via OCT4[10]. Here, we confirmed that E2F6 and E2F3 are highly expressed in liver cancer and liver TICs, and are the top two E2F TFs expressed in liver tumorigenesis. *E2F6* and *E2F3* knockout cells harbor weak self-renewal and tumorigenesis capabilities. Notably, E2F antagonist HLM006474 inhibited the expansion and metastasis of liver cancer, as well as the self-renewal and tumor initiation of liver TICs. This suggests that E2F6 and E2F3 are potential therapeutic targets for liver cancer and elimination of liver TICs. We also provide preliminary evidence showing that E2F6 and E2F3 mediate their effects via the Wnt/β-catenin signaling pathway, although the specific molecular mechanisms need to be studied in more detail.

In summary, we have identified a functional rt-circRNA, rtcisE2F, which is required for the self-renewal of liver TICs. rtcisE2F-dependant E2F6/E2F3 may be useful for determining the diagnosis and prognosis of liver cancer, and might be used as targets for eradicating liver TICs.

## Materials and Methods

### Reagents and Antibodies

Anti-CD133 (catalog no. 130-090-853) and PE conjugated anti-CD133 antibodies were purchased from Miltenyi Biotec. YTHDF2 (catalog no. 80014), SOX2 (catalog no. 23064S), AXIN2 (catalog no. 20540-1-AP) and c-MYC (catalog no. 2276S) antibodies were from Cell Signaling Technology. Anti-E2F3 (catalog no. 27615-1-AP) and IGF2BP2 (catalog no. 11601-1-AP) antibodies were from Proteintech. Anti-digoxin (catalog no. ab51949) antibody was obtained from Abcam. β-actin (catalog no. RM2001) antibody was purchased from Beijing Ray antibody Biotech. ZIC2 (catalog no. ARP35821_P050) antibody was purchased from Aviva Systems Biology. Alexa-594-, Alexa-488- and Alexa-647-conjugated anti-rabbit and anti-mouse secondary antibodies were purchased from Invitrogen. HRP-conjugated secondary antibody was purchased from Sungene Biotech. Polymer HRP and AP detection kits were from GBI labs. Biotin labeld RNA mix (catalog no. 11685597910) was from Rhoche. Chemiluminescent nucleic acid detection module (catalog no. 89880) was from Thermo Scientific. ChIP assay kit (catalog no. 17-295) was from Miltenyi Biotec. Supplements N2 and B27 were purchased from Life Technologies.

### Isolation of liver cancer cells and TICs

Primary liver cancer cells were obtained from liver cancer patients. Fresh liver cancer tissues were washed two or three times and kept in DMEM/F12 medium supplemented with 1000 U/ml penicillin and 1000 U/ml streptomycin, and transferred to lab on ice quickly. Samples were then washed with pre-cooled sterile PBS supplemented with 100 U/ml penicillin and 100 U/ml streptomycin, cut into small fragments, and digested in HBSS containing 0.03% pronase, 0.05% type IV collagenase, and 0.01% deoxyribonuclease for 30 min at 37 °C. Then sample was filtered through 100 μm nylon filter, centrifuged for 2 min at 50 x g in 4 °C and HCC primary cells were in precipitation.

Liver TICs are obtained from FACS sorting or sphere formation assay. For TIC enrichment, liver cancer cells were stained with CD133 antibody, and CD133^+^ liver TICs were enriched by FACS. For sphere formation, 5000 primary cells were seeded into Ultra Low Attachment 6-well plates (Corning Incorporated Life Sciences), and cultured in DMEM/F12 (Life Technologies) supplemented with N2, B27, 20 ng/ml EGF and 20 ng/ml bFGF (Millipore).

### CRISPR-Cas9 knockout and CRISPRi

*E2F6*, *E2F3*, *IGF2BP2* and *YTHDF2* knockout cells were generated by standard approach, with minor modifications[55]. Generally, sgRNAs were designed and cloned into LentiCRISPRv2 (Puro, catalog no. 52961). LentiCRISPRv2, pVSVg (catalog no. 8454) and psPAX2 (catalog no. 12260) were used to generate CRISPR-Cas9 lentivirus for liver TIC infection.

For CRISPRi, sgRNAs were designed and cloned. dCas9-KRAB (catalog no. 46911), lentivirus and sgRNA lentivirus were packaged for liver TIC infection. All plasmids were obtained from Addgene.

The sequences of sgRNAs used for CRISPR-Cas9 and CRISPRi were listed in Supplementary Table 1.

### Tumor initiation assay

For tumor initiation assay, 10, 1×10^2^, 1×10^3^, 1×10^4^, and 1×10^5^ *rtcisE2F* knockout, overexpression and control cells were subcutaneously injected into 6-week-old BALB/c nude mice. Tumor initiation was detected after 3 months and the ratios of tumor-free mice were calculated.

### Transwell assay

Transwell assay was performed as described[56]. Briefly, 1×10^5^ *rtcisE2F* knockout, overexpression, E2F inhibitor HLM006474 treated and control cells were seeded into a top chamber with matrigel-coated membrane (ThermoFisher Scientific) without FBS, and then medium supplemented with FBS is added into lower chamber as a chemoattractant. 36 hours later, the cells on the lower surface of chamber membrane were fixed with methanol, stained with crystal violet, and finally imaged with Nikon Eclipse Ti2-U.

### Tagged RNA affinity purification

Tagged RNA affinity purification (TRAP) assay was performed as described[41]. Briefly, rtcisE2F conjugated with MS2 sequence and GST conjugated MCP (MS2 coat protein), which was cloned from plasmid (Addgene no. 75384), were overexpressed in liver cancer cells. rtcisE2F binding proteins were enriched through GST pulldown assay, and identified by Silver staining and mass spectrum, or detected by Western blot with IGF2BP2 and YTHDF2 antibodies.

### *In situ* hybridization

Digoxin-conjugated rtcisE2F probes were designed according to protocols of Biosearch Technologies, and *in situ* hybridization was performed under non-denatured conditions as described (https://www.biosearchtech.com/). The samples were visualized with DAB (diaminobezidin), counterstained with hematoxylin, dehydrated and mounted. Finally, samples were observed with Thermo CX7 LZR confocal microscope.

### Western blot

Western blot is performed according to the standard method as previously described[57]. Simply, liver cancer cells and TICs are crushed with 1×SDS-loading buffer, and boiled 15 minutes, followed by electrophoresis. Protein samples was then transferred to nitrate cellulose (NC) membrane, incubated with primary antibodies overnight at 4℃, and then incubated with HRP-labeled secondary antibodies, finally the HRP signals were visualized by ultra-sensitive enhanced chemiluminescent (ECL) substrate.

### Chromatin Isolation by RNA purification (ChIRP)

ChIRP assay was performed as described previously[58]. Briefly, spheres were cross-linked with 1% glutaraldehyde and lyzed with lysis buffer, followed by sonication to get 200-500 bp DNA fragments (Bioruptor). Digoxin labelled rtcisE2F and control probes were added into sphere lysates for 4 hour incubation at 37°C, and then digoxin antibody was used for the enrichment of chromatin components. The enrichment of *E2F6* and *E2F3* promoter was detected by real-time PCR, and the primers were in Supplementary Table 2.

### RNA immunoprecipitation

*rtcisE2F* knockout, overexpressing and control spheres were lyzed in RNase-free RIPA buffer (150 mM NaCl, 0.5% sodium deoxycholate, 0.1% SDS, 1% NP-40, 1 mM EDTA and 50 mM Tris, pH 8.0, containing protease-inhibitor cocktail and RNase inhibitor), followed by ultrasonication. Samples were centrifuged and supernatants were collected for preclear with Protein A/G. YTHDF2 and IGF2BP2 antibodies were incubated with Protein A/G, and then added to sphere lysates for 4 hour incubation. Total RNA in eluate was extracted and *E2F6*/*E2F3* mRNA enrichment was detected through real-time PCR, and the primers were in Supplementary Table 2.

### Signaling pathway activity reporter system

Signaling pathway (including Wnt/β-catenin, Notch, Hedgehog, mTOR, NF-kB, P38, JNK, ERK and PKA) reporter plasmids and control plasmids were overexpressed in HLM006474 treated and control cells, and the activity of signaling pathway can be detected by FACS. For example, Wnt activity= (TOP-GFP intensity)/(FOP-GFP intensity). The plasmids used in this assay are: TOP-GFP (addgene no. 35489), FOP-GFP (addgene no. 35490), 12XCSL-d1EGFP (addgene no. 47684), 7Gli:GFP (addgene no.110494), TORCAR (addgene no. 64927), TORCAR(T/A) (addgene no. 64928), NF-kB-eGFP (addgene no. 118093), p38KTRmCerulean3 (addgene no. 59155), JNKKTRmRuby2 (addgene no. 59154), ERKKTRClover (addgene no. 59150), PKAKTRClover (addgene no.59153).

### RNA stability detection

*rtcisE2F* knockout, *YTHDF2* knockout, *IGF2BP2* knockout and control cells were treated with 2 μg/ml actinomycin D (Act-D) to block transcription, and equant cells were obtained at time points of 0, 0.5, 1, 2, 3 and 4 hours after Act-D treatment. *E2F6* and *E2F3* mRNA levels were detected by real-time PCR or Northern blot, and normalized with E2F6/E2F3 levels at 0 hour. The primers were in Supplementary Table 2.

## Data viability

All data supporting the findings of this study are available from the corresponding author on reasonable request.

## Abbreviations

circRNA: circular RNA
HCC: Hepatocellular carcinoma
TIC: Tumor initiating cell
WT: Wild type
KO: Knockout
oe: overexpression
RIP: RNA immunoprecipitation
TRAP: Tagged RNA affinity purification
ISH: In Situ Hybridization
FISH: Flourescence in situ hybridization

## Declarations

### Ethics approval and consent to participate

Not applicable.

### Consent for publication

All authors consent to the publication of the manuscript in Molecular Cancer.

### Authors’ information

Zhenzhen Chen: School of Life Sciences, Zhengzhou University, Zhengzhou 450001, China;

Benyu Liu: The Academy of Medical Science, College of Medical, Zhengzhou University, Zhengzhou, Henan, China;

Lan Huang: Biotherapy Center, the First Affiliated Hospital of Zhengzhou University, 1st Jianshe East Road, Zhengzhou, Henan 450052, China;

Xiang Zhong: School of Basic Medical Sciences, Henan University, Kaifeng, 475004, China;

Zhongyi Yan: School of Basic Medical Sciences, Henan University, Kaifeng, 475004, China;

Pingping Zhu: School of Life Sciences, Zhengzhou University, Zhengzhou 450001, China;

## Acknowledgements

This work was supported by Strategic Priority Research Programs of the Chinese Academy of Sciences (2020YFA0803500), National Natural Science Foundation of China (31922024, 81872411, 31771638), Program for Innovative Talents of Science and Technology in Henan Province (18HASTIT042), Young Talent Support Project from Chinese Association of Science and Technology (YESS20170042), Science Foundation for Excellent Young Scholars in Henan (202300410358), Young Talent Support Project from Henan province (2018HYTP002). We thank the supporting grants from Zhengzhou University to Pingping Zhu, and the technical support from Modern Analysis and Computer Center of Zhengzhou University.

## Author contributions

Z.C. and B.L. designed and performed experiments, analyzed data and wrote the paper; L.H provided HCC samples, performed experiments, analyzed data and wrote the paper; X.Z., and Z.Y. performed experiments and analyzed data, P.Z. initiated the study, designed and performed experiments, analyzed data and wrote the paper.

