## Supplemental Figures and Tables for "rtcisE2F drives liver TIC self-renewal and metastasis via m^6^A-modulated mRNA stability of *E2F6* and *E2F3*"

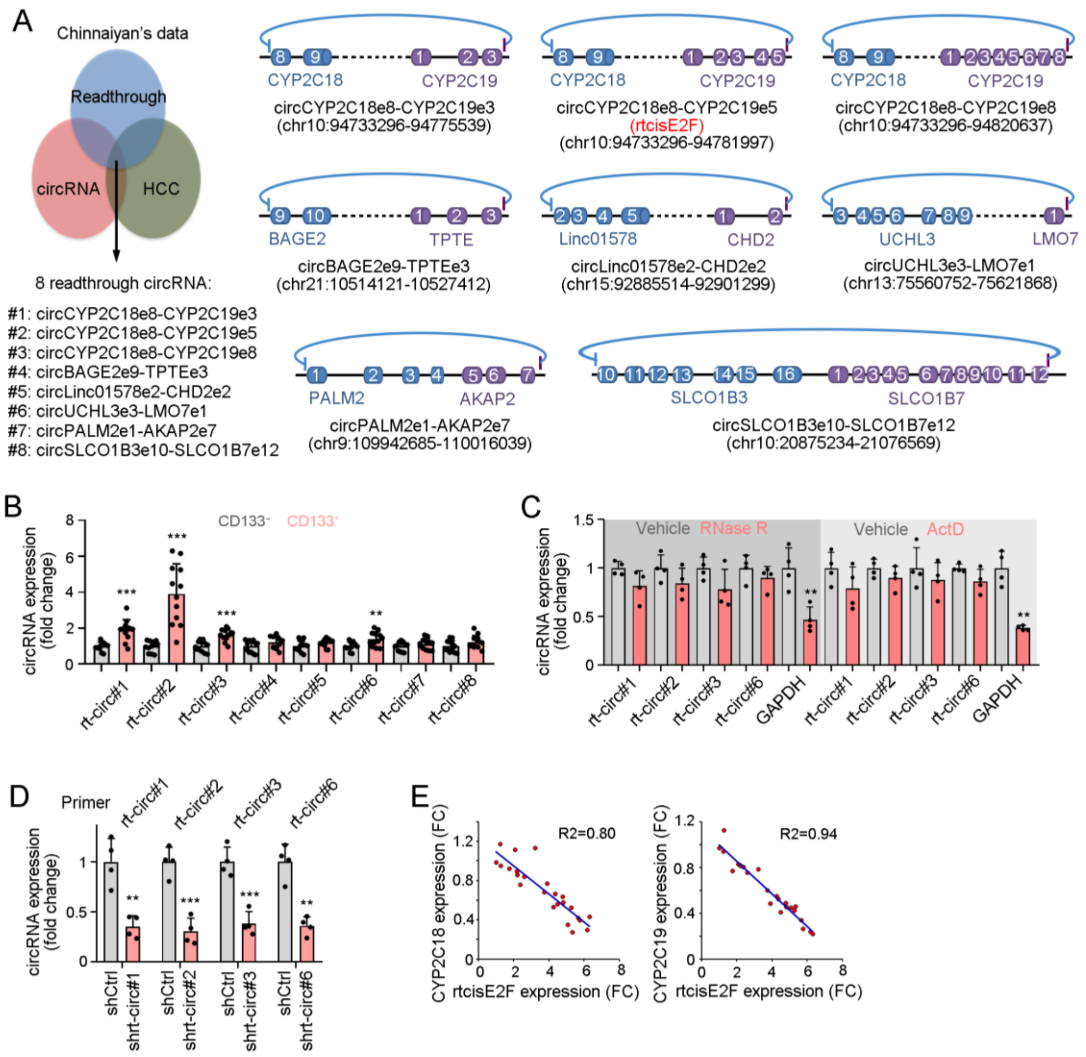


**Supplementary Figure 1. Identification and validation of rt-circRNAs in liver TICs.** (A) Schematic diagram of 8 rt-circRNAs expressed in liver cancer. circRNA screening process was shown in the left panel, and genomic compositions of circRNAs were depicted in the right panel. circCYP2C18e8-CYP2C19e5 is rtcisE2F. (B) Quantitative real-time PCR analysis for indicated rt-circRNA expression in $CD133^-$ and $CD133^+$ cells isolated from liver cancer samples. All expression levels were normalized to those in $CD133^-$ cells. (C) Validation of circRNAs using RNase R treatment and Actinomycin D (ActD). Left, total RNAs extracted from liver TICs were treated with or without 3 U/g RNase R for 1 h. Right, liver TICs were treated with or without 2 μg/ml actinomycin D (ActD), followed by RNA extraction and real-time PCR analysis of rt-circRNAs. (D) Quantitative real-time PCR to analyze rt-circRNA knockdown efficiency in indicated rt-circRNA silenced cells. (E) Co-expression of rtcisE2F and its parent genes CYP2C18 and CYP2C19. All expression levels were normalized to those in peri-tumor. For C and D, *n* = 4 independent experiments.

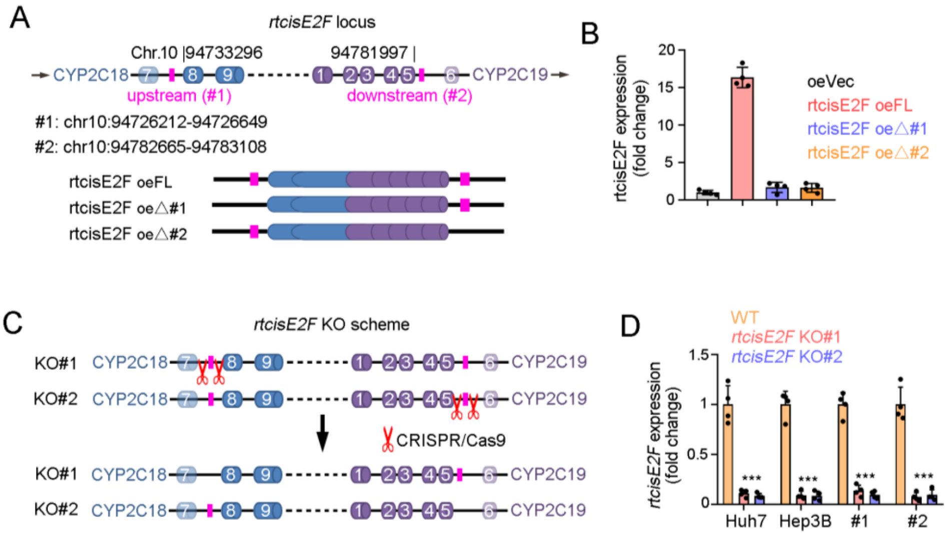


**Supplementary Figure 2. Generation of *rtcisE2F* knockout cells.** (A, B) Minigene assay for the necessity of upstream and downstream reverse complementary sequences in rtcisE2F biogenesis (A), and rtcisE2F expression was detected through quantitative real-time PCR (B). FL, full length, △1, full length without #1 sequence (upstream sequence), △2, full length without #2 sequence (downstream sequence). (C) Schematic diagram of CRISPR-Cas9 based *rtcisE2F* knockout strategy. Upstream and downstream reverse complementary sequences were deleted in KO#1 and KO#2 cells, respectively. (D) Quantitative real-time PCR analysis to confirm knockout efficiency. Data are shown as mean ± s.d. **P* < 0.05; ***P* < 0.01; ****P* < 0.001, by one-tailed Student's T-test.

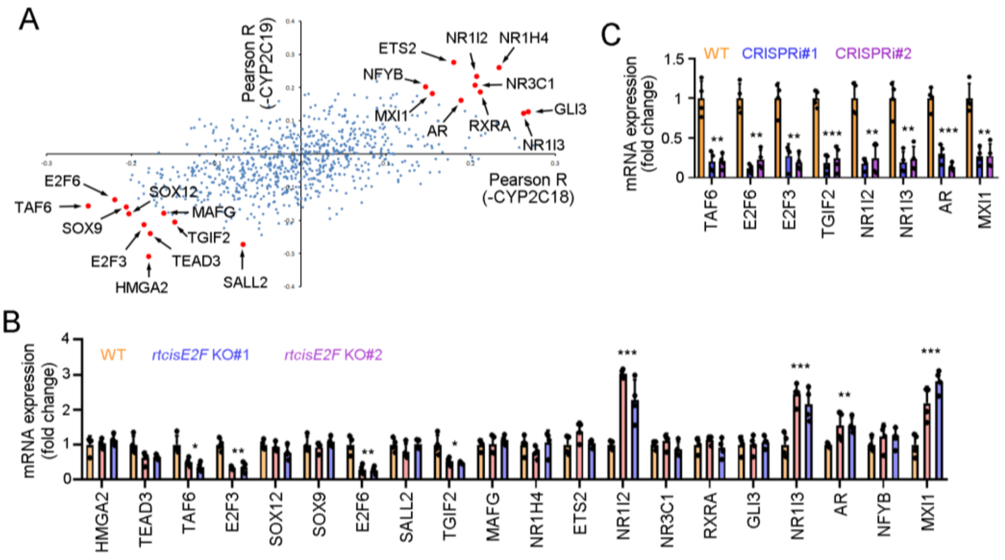


**Supplementary Figure 3. Identification of rtcisE2F target genes.** (A) Co-expression of transcription factors with rtcisE2F parent genes CYP2C18 and CYP2C19, showing top10 positively correlated and negatively correlated genes. TF expression levels were analyzed from 224 liver cancer patients, according to online-available dataset GSE14520. (B) Real-time PCR to detect the expression levels of indicated TFs in *rtcisE2F* knockout and WT primary cells. All expression levels were normalized to those in WT cells. (C) Quantitative real-time PCR for TF depletion efficiency via CRISPRi-KRAB strategy. In all panels, data are shown as mean ± s.d. *$P$ < 0.05; **$P$ < 0.01; ***$P$ < 0.001, by one-tailed Student’s T-test.

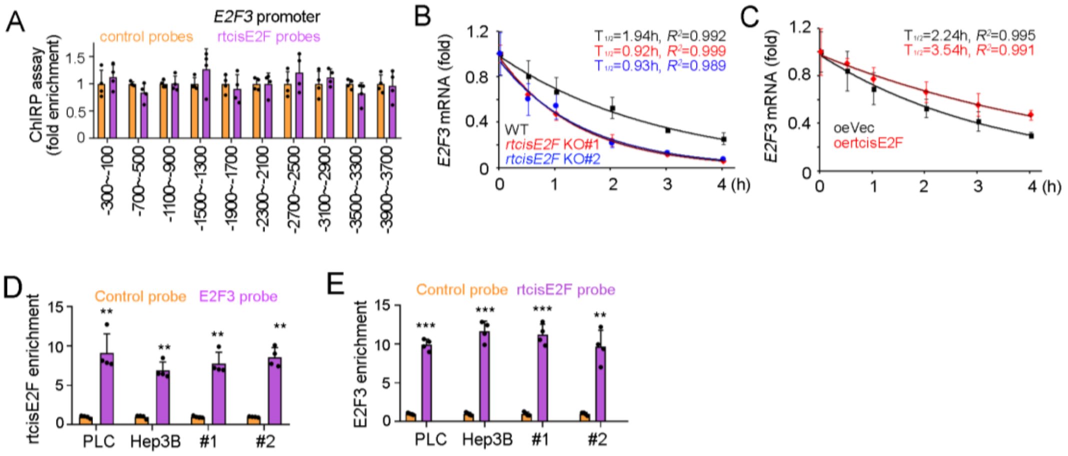


**Supplementary Figure 4. rtcisE2F promotes *E2F3* mRNA stability.** (A) Quantitative real-time PCR for the enrichment of *E2F3* promoter in eluate of ChIRP assay with rtcisE2F probes. (B, C) Quantitative real-time PCR analysis of *E2F3* mRNA stability in *rtcisE2F* knockout TICs (B) or overexpressing TICs (C) treated with 2 μg/ml actinomycin D for the indicated times. (D, E) Quantitative real-time PCR analysis of RNA pulldown eluate for the interaction of *E2F3* mRNA and rtcisE2F. Data are shown as mean ± s.d. ***P* < 0.01; ****P* < 0.001, by one-tailed Student's T-test.

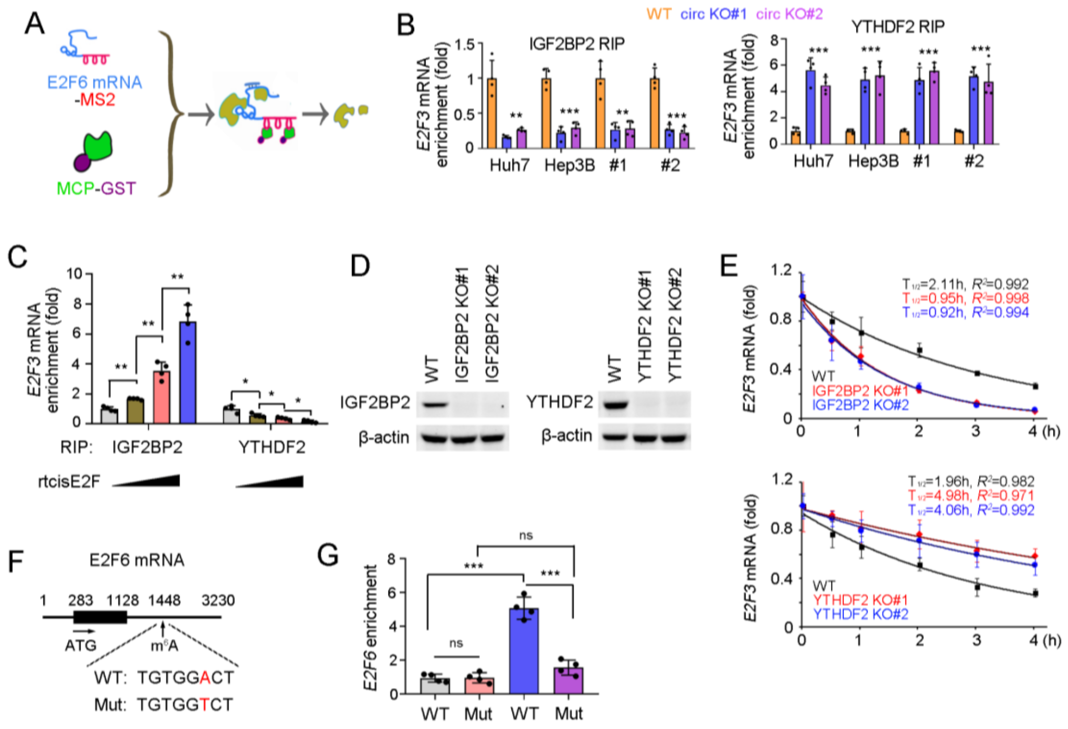


**Supplementary Figure 5. rtcisE2F is involved in the association of *E2F3* mRNA and m$^6$A readers IGF2BP2/YTHDF2.** (A) Schematic diagram of TRAP assay, for which MS2 conjugated *E2F6* mRNA and MCP-GST were used. *E2F6* mRNA binding proteins were visualized via silver staining and identified via mass spectrum, and confirmed by Western blot. (B, C) Quantitative real-time PCR of *E2F3* mRNA in RIP eluate using IGF2BP2 antibody (left) and YTHDF2 antibody (right). For B, *rtcisE2F* knockout and WT cells were used. For C, gradient rtcisE2F transcripts were added into cell lysate. (D) Western blot to confirm the KO efficiency of IGF2BP2 and YTHDF2, which were generated through CRISPR-Cas9 approach. (E) Quantitative real-time PCR analysis of *E2F3* mRNA in *IGF2BP2* knockout cells (upper panel) and *YTHDF2* knockout cells (lower panel), treated with 2 μg/ml actinomycin D for indicated time points. (F) Schematic diagram of E2F6 mutation, which harbors no m$^6$A site. (G) Quantitative real-time PCR analysis of *E2F6* mRNA in m$^6$A RNA immunoprecipitation, in which E2F6 WT and mutant (Mut) spheres were used. In all panels, data are shown as mean ± s.d. **$P < 0.01$; ***$P < 0.001$, by one-tailed Student's T-test. For all representative images, at least three independent experiments were performed with similar results.

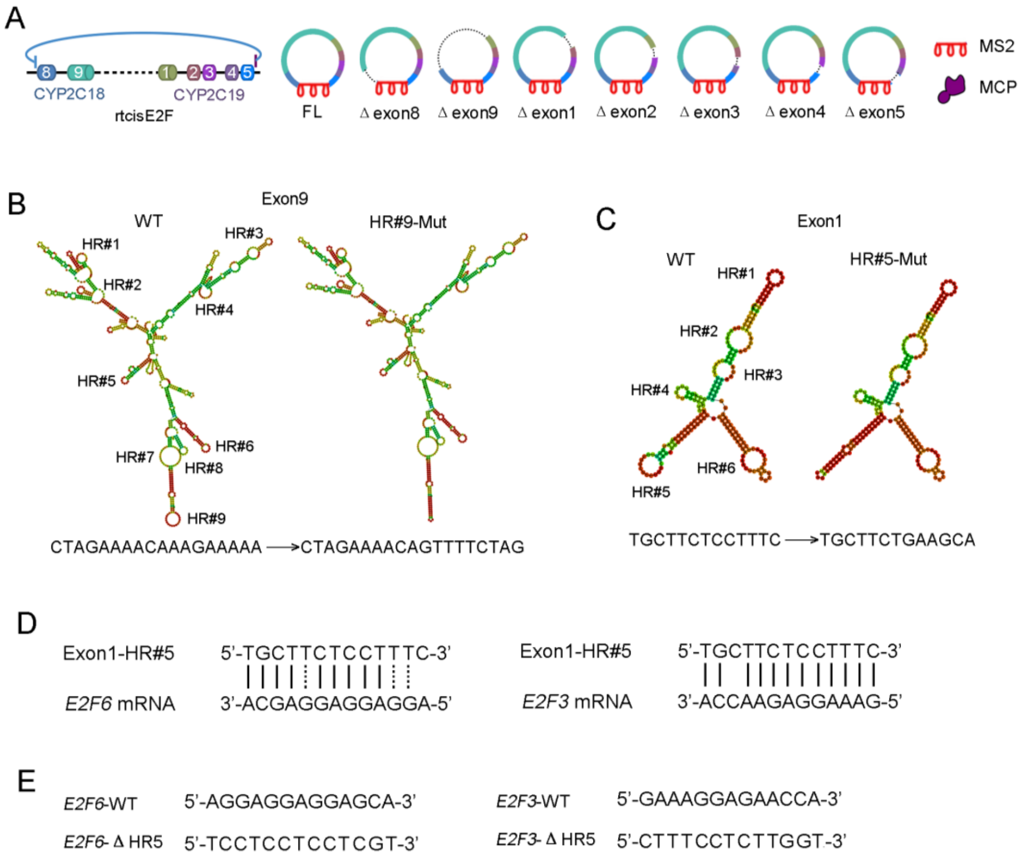


**Supplementary Figure 6. rtcisE2F promotes the interaction of IGF2BP2 and *E2F6/E2F3* mRNA as a scaffold.** (A) Schematic diagram of truncate rtcisE2F. The seven exons were deleted individually. (B, C) Prediction structure of the second exon (B) and third exon (C) of rtcisE2F. Structure prediction was based on minimum free energy (MFE). HR: hairpin loop region. (D) Diagram of *rtcisE2F* HR#5 sequence was shown in upper panel and matching sequences in *E2F6* and *E2F3* mRNAs were shown in lower panel. (E) Schematic diagram of *E2F6* and *E2F3* mutation, which lost rtcisE2F-bindng capacity.

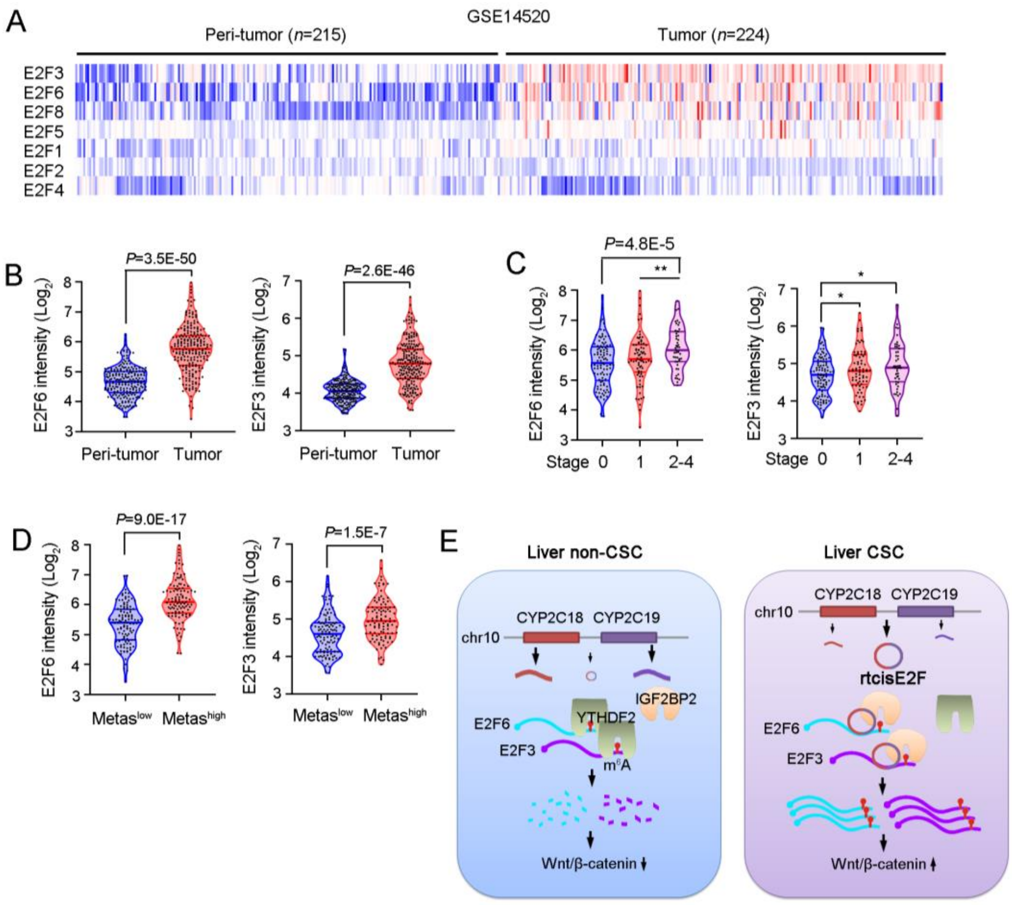


**Supplementary Figure 7. E2F6 and E2F3 are highly expressed in liver cancer, especially in advanced liver cancer.** (A) Heatmap showing the expression landscape of E2F TF family, among which E2F3 and E2F6 were most significantly up-regulated along liver tumorigenesis. (B-D) Violin plots of E2F6 and E2F3 expression in liver tumors and peri-tumors (B), liver tumors with different clinical stages (C) or different metastasis capacity (D). Individual samples are shown as black dots, with medium, minimum, maximum and quarter levels shown as lines. (E) The molecular mechanism of rtcisE2F. rt-circRNA rtcisE2F is originated from two adjacent genes CYP2C18 and CYP2C19, involves in the binding between $m^6A$ *E2F6*/*E2F3* mRNAs and $m^6A$ readers IGF2BP2/YTHDF2, and finally promotes the stability of *E2F6*/*E2F3* mRNAs. E2F6 and E2F3 drive the activation of Wnt/β-catenin signaling and the self-renewal of liver TICs.

### Supplementary Table 1. sgRNA sequences used in this study

| sgRNA | Sequences |
|---|---|
| *rtcisE2F* KO#1 (up) | 5'-AATGTGCAGTTTTGTTACAT-3' |
| *rtcisE2F* KO#1(down) | 5'-TGTGTTTCGTATGTTTATTG-3' |
| *rtcisE2F* KO#2 (up) | 5'-TATTCACAATAGCCAAGACT-3' |
| *rtcisE2F* KO#2(down) | 5'-GATACATGTGCAGAACGTGC-3' |
| *IGF2BP2* KO#1 | 5'-TGCATATGTGACGTTGACAA-3' |
| *IGF2BP2* KO#2 | 5'-GTGGGGACCAGGATCCGCAG-3' |
| *YTHDF2* KO#1 | 5'-GCATGAATACTATAGACCAA-3' |
| *YTHDF2* KO#2 | 5'-GCTGAGAAGTCAATCCCACT-3' |
| *E2F6* KO#1 | 5'-TCTTGGATTTCTTTTCAACG-3' |
| *E2F6* KO#2 | 5'-CCAACAAAAGAAGCTACAGG-3' |
| *E2F3* KO#1 | 5'-TCATACCGCGTTTTTTCTGA-3' |
| *E2F3* KO#2 | 5'-GGGTTTTTAAACCATCTGAG-3' |
| TAF6 (CRISPRi) | 5'-CAGGACCTGCTCCCACGAGG-3' |
| E2F6 (CRISPRi) | 5'-GCCGGGAGAGGATCGGTGCG-3' |
| E2F3 (CRISPRi) | 5'-GGGATACGGTTTACGCGCCA-3' |
| TGIF2 (CRISPRi) | 5'-GGATCTGACGTCAGGCCGCG-3' |
| NR1I2 (CRISPRi) | 5'-ACAAGATTGTCTCATATCCG-3' |
| NR1I3 (CRISPRi) | 5'-GGGCACAGACCCTGGACTCA-3' |
| AR (CRISPRi) | 5'-AAATGCAACAGTTTGCGAGT-3' |
| MXI1 (CRISPRi) | 5'-GGGAGATTGAATGAACCTAC-3' |

**Supplementary Table 2. PCR primers used in this study**

| Primers | Sequences |
|---|---|
| 18S (Forward) | 5'-AACCCGTTGAACCCCATT-3' |
| 18S (Reverse) | 5'-CCATCCAATCGGTAGTAGCG-3' |
| actin (Forward) | 5'-GGCTGTATTCCCCTCCATCG-3' |
| actin (Reverse) | 5'-CCAGTTGGTAACAATGCCATGT-3' |
| rt-circ#1 (Divergent Forward) | 5'-CAGCAATGGAAAGAGATGGA-3' |
| rt-circ#1 (Divergent Reverse) | 5'-TTCTTAAAGTTGCCACTCTT-3' |
| rt-circ#2 (Divergent Forward) | 5'-CAAGAATCGATGGACATCAA-3' |
| rt-circ#2 (Divergent Reverse) | 5'- TTCTTAAAGTTGCCACTCTT -3' |
| rt-circ#3 (Divergent Forward) | 5'- AAGAATTTCCCAACCCAGAG -3' |
| rt-circ#3 (Divergent Reverse) | 5'- TTCTTAAAGTTGCCACTCTT -3' |
| rt-circ#4 (Divergent Forward) | 5'- AGTTATGGATTCACTACCA -3' |
| rt-circ#4 (Divergent Reverse) | 5'- CACCATCTGT TCCTTCCACT -3' |
| rt-circ#5 (Divergent Forward) | 5'- AGATGATGAG AAATAAGGAC -3' |
| rt-circ#5 (Divergent Reverse) | 5'- AATCTGTGTT TGCTTTGAAG -3' |
| rt-circ#6 (Divergent Forward) | 5'- GAGGAGGCAG AGGCCAACTG -3' |
| rt-circ#6 (Divergent Reverse) | 5'- GAAGTAAGAC TGCACAGACT -3' |
| rt-circ#7 (Divergent Forward) | 5'- GAATGTGCTG CTAAAGGAAG -3' |
| rt-circ#7 (Divergent Reverse) | 5'- GCACACAGGC AGGGGAAAGA -3' |
| rt-circ#8 (Divergent Forward) | 5'- GGAGCACGAG GGGCTTGTAG -3' |
| rt-circ#8 (Divergent Reverse) | 5'- GCTTACTTGT AACAATGTCA -3' |
| E2F6 (Forward) | 5'- TCCATGAACAGATCGTCATTGC-3' |
| E2F6 (Reverse) | 5'- TCCGTTGGTGCTCCTTATGTG-3' |
| E2F3 (Forward) | 5'- AGAAAGCGGTCATCAGTACCT-3' |
| E2F3 (Reverse) | 5'- TGGACTTCGTAGTGCAGCTCT-3' |
| TAF6 (Forward) | 5'- CTAACGGATGAGGTCAGCTACC-3' |
| TAF6 (Reverse) | 5'- AAGGCGTAGTCAATGTCACTG-3' |
| TGIF2 (Forward) | 5'- ATGTCGGACAGCGATCTAGG-3' |

| | |
|---|---|
| TGIF2 (Reverse) | 5’- TCCCGGAGGATCTTTACTGAC-3’ |
| NR1I2 (Forward) | 5’- GATGGAGGTCTTCAAATCTGCC-3’ |
| NR1I2 (Reverse) | 5’- GGCCCTTCTGAAAAACCCCT-3’ |
| NR1I3 (Forward) | 5’- ATATGGGCCGAGGAACTGTGT-3’ |
| NR1I3 (Reverse) | 5’- GGCGTGGAAATGATAGCCTGT-3’ |
| AR (Forward) | 5’- CTGGGAAGGGTCTACCCAC-3’ |
| AR (Reverse) | 5’- GGTGCTATGTTAGCGGCCTC-3’ |
| MXI1 (Forward) | 5’- GCGCCTTTGTTTAGAACGCTT-3’ |
| MXI1 (Reverse) | 5’- AATGCTGTCCATTCGTATTCGT-3’ |
| GAPDH (Forward) | 5’- AGGTCGGTGTGAACGGATTTG-3’ |
| GAPDH (Reverse) | 5’- TGTAGACCATGTAGTTGAGGTCA-3’ |
| HMGA2 (Forward) | 5’- GAGCCCTCTCCTAAGAGACCC-3’ |
| HMGA2 (Reverse) | 5’- TTGGCCGTTTTTCTCCAATGG-3’ |
| TEAD3 (Forward) | 5’- CAACCAGCACAATAGCGTCCA-3’ |
| TEAD3 (Reverse) | 5’- CTGAAAGCTCTGCTCGATGTC-3’ |
| SOX12 (Forward) | 5’- AAGAGGCCGATGAACGCATT-3’ |
| SOX12 (Reverse) | 5’- TAGTCCGGGTAATCCGCCAT-3’ |
| SOX9 (Forward) | 5’- GAGCCGGATCTGAAGAGGGA-3’ |
| SOX9 (Reverse) | 5’- GCTTGACGTGTGGCTTGTTC-3’ |
| SALL2 (Forward) | 5’- CTCGCTCACCAGAACTCATGT-3’ |
| SALL2 (Reverse) | 5’- GGGGATTGCTGTGCTCTGTA-3’ |
| MAFG (Forward) | 5’- ATGACGACCCCCAATAAAGGA-3’ |
| MAFG (Reverse) | 5’- CACCGACATGGTTACCAGC-3’ |
| NR1H4 (Forward) | 5’- GCTTGATGTGCTACAAAAGCTG-3’ |
| NR1H4 (Reverse) | 5’- CGTGGTGATGGTTGAATGTCC-3’ |
| ETS2 (Forward) | 5’- CCCCTGTGGCTAACAGTTACA -3’ |
| ETS2 (Reverse) | 5’- AGGTAGCTTTTAAGGCTTGACTC -3’ |
| NR3C1 (Forward) | 5’- AGCTCCCCCTGGTAGAGAC -3’ |
| NR3C1 (Reverse) | 5’- GGTGAAGACGCAGAAACCTTG -3’ |

| | |
|---|---|
| RXRA (Forward) | 5’- ATGGACACCAAACATTTCCTGC-3’ |
| RXRA (Reverse) | 5’-CCAGTGGAGAGCCGATTCC-3’ |
| GLI3 (Forward) | 5’-CACAGCTCTACGGCGACTG-3’ |
| GLI3 (Reverse) | 5’-CTGCATAGTGATTGCGTTTCTTC-3’ |
| NFYB (Forward) | 5’-GCCTCCCAGCTAGGGATTTC -3’ |
| NFYB (Reverse) | 5’-TTCCTGTTTGAGGTATGGCATTT-3’ |
| E2F3promt-300 (Forward) | 5’-GGAGAGAAGGAAATTCTTTG-3’ |
| E2F3promt-100 (Reverse) | 5’-GGACCGTTCCCACAGCCCAT-3’ |
| E2F3promt-700 (Forward) | 5’-TGAACCTTCTGCCCCCACGA-3’ |
| E2F3promt-500 (Reverse) | 5’-CGCAGGAGATTTTTCCAGAT-3’ |
| E2F3promt-1100 (Forward) | 5’-TTTCTAGTTTACTTTTTAAA-3’ |
| E2F3promt-900 (Reverse) | 5’-ATCCGCCAAGCAGTAATCCG-3’ |
| E2F3promt-1500 (Forward) | 5’-GAGCCTGGTGCAAGGTTTAA-3’ |
| E2F3promt-1300 (Reverse) | 5’-AGGTGATGAGGATTGCAACA-3’ |
| E2F3promt-1900 (Forward) | 5’-TGCCCTAAAGATAGTGTCAC-3’ |
| E2F3promt-1700 (Reverse) | 5’-TGAGTAGCTGGGACTACAGG-3’ |
| E2F3promt-2300 (Forward) | 5’-ATAAATATCTGAAAGCTTTC-3’ |
| E2F3promt-2100 (Reverse) | 5’-GGTGCAACATACACAGTAAT-3’ |
| E2F3promt-2700 (Forward) | 5’-CAGCCTAATATTTGTATTTT-3’ |
| E2F3promt-2500 (Reverse) | 5’-AGCTTTGGAGTCAGAGTGAT-3’ |
| E2F3promt-3100 (Forward) | 5’-TGCTATATGCCTGGCACTTT-3’ |
| E2F3promt-2900 (Reverse) | 5’-CTGGGAGGCGGAGTTGCAGT-3’ |
| E2F3promt-3500 (Forward) | 5’-AGAAGTCTTGGGGCAGTCAG-3’ |
| E2F3promt-3300 (Reverse) | 5’-ACTCAGAATAATAACATAAA-3’ |
| E2F3promt-3900 (Forward) | 5’- GGAGGCTGGAAGTCCAAAAT -3’ |
| E2F3promt-3700 (Reverse) | 5’-AG GGGACACCCAGAGGAGGC-3’ |
| E2F6promt-300 (Forward) | 5’-CTGCACTCCAGCCTGGGCGAC-3’ |
| E2F6promt-100 (Reverse) | 5’- GTCGCCCAGGCTGGAGTGC -3’ |
| E2F6promt-700 (Forward) | 5’-GGAAGGAAGGAAGGAAGGAA-3’ |

| | |
|---|---|
| E2F6promt-500 (Reverse) | 5'- ACCTTGGCCTCTCAAGTGCT -3' |
| E2F6promt-1100 (Forward) | 5'-ATACCACGGTGTGTTGCCTT-3' |
| E2F6promt-900 (Reverse) | 5'- ACCTTGGCCTCTCAAGTGCT-3' |
| E2F6promt-1500 (Forward) | 5'-TAGAAGCCAGGCATGGGTGT-3' |
| E2F6promt-1300 (Reverse) | 5'- CAAACTCCTGGCCTCAAGTG-3' |
| E2F6promt-1900 (Forward) | 5'-GAATCTTCCTAAAGATCTGA-3' |
| E2F6promt-1700 (Reverse) | 5'- GCATTCCACATGCATTTGCT-3' |
| E2F6promt-2300 (Forward) | 5'-CTTCCTCTGCTGTGCTGCTA-3' |
| E2F6promt-2100 (Reverse) | 5'- AAGCATCTCTCATGGGCCA-3' |
| E2F6promt-2700 (Forward) | 5'-CTGAGCCTCTGTACCCTCAT-3' |
| E2F6promt-2500 (Reverse) | 5'- TACAAAAATTAGCCGGGCGT-3' |
| E2F6promt-3100 (Forward) | 5'-GGTCATTTCCTCACGGACCT-3' |
| E2F6promt-2900 (Reverse) | 5'- AATGGGGCATGGGCTTTGGG-3' |
| E2F6promt-3500 (Forward) | 5'-GTTCTCAGCTTGGCTCTTCC-3' |
| E2F6promt-3300 (Reverse) | 5'- AAACTAAAATTGTAGATATC-3' |
| E2F6promt-3900 (Forward) | 5'-TAAGAGCTAAGTGGTTGCAT-3' |
| E2F6promt-3700 (Reverse) | 5'- AAAG GACAGAACTCAGTCAT-3' |
